# A cytochrome P450 involved in apocarotenoid signaling enhances plant photosynthetic capacity and photooxidative stress tolerance

**DOI:** 10.1101/2024.12.20.629731

**Authors:** Madhu Tiwari, Brigitte Ksas, Betrand Légeret, Stefano Caffarri, Michel Havaux

## Abstract

β-cyclocitric acid and its precursor β-cyclocitral are signaling apocarotenoids which trigger defense and detoxification mechanisms enhancing plant tolerance to abiotic stresses. From a transcriptomic analysis of Arabidopsis plants exposed to each apocarotenoid over several exposure times, we identified a gene (*CYP81D11*) encoding a cytochrome P450 that is strongly induced under all conditions and is under the control of the TGAII-SCL14 transcription regulator. Overexpressing the *CYP81D11* gene in Arabidopsis led to a very tolerant phenotype to high light stress while a *CYP81D11*- deficient mutant was photosensitive. The transcriptomic profile of the *CYP81D11* overexpressor revealed a selective upregulation of genes related to photosynthesis and the response to high light stress. Photosynthetic electron transport, CO_2_ fixation and biomass production were noticeably improved by high expression levels of *CYP81D11*. These effects occurred in high light, not in low light, and were associated with a noticeable reduction of singlet oxygen photoproduction. These findings indicate that CYP81D11 is a key component of β-cyclocitral-induced stress tolerance, acting through enhancement of the photosynthetic capacity of leaves. This gene could be a new target for improving photosynthesis, particularly in sunny environments.

---

Plants often encounter abiotic constraints in the wild, such as drought, extreme temperatures or high light intensity, which pose serious challenges for maintaining cellular stability and photosynthetic efficiency (Apel and Hirt 2004, Li et al. 2009, Ort et al. 2015). For instance, high light intensity can exceed the photosynthetic capacity of chloroplasts, resulting in the absorption of more light energy than can be utilized in photochemical reactions (Ort 2001). This surplus energy creates a state of overexcitation in the photosynthetic apparatus, leading to the formation of reactive oxygen species (ROS). The major photodamaging ROS in plant leaves is singlet oxygen (^1^O_2_) (Triantaphylides et al. 2008) which results from the interaction of triplet excited chlorophyll molecules with molecular oxygen (Triantaphylidès and Havaux, 2009). On the other hand, by inducing stomatal closure and hence limiting CO_2_ supply to the chloroplasts, drought stress inhibits photosynthesis, also leading to excess light energy and ROS production (Smirnoff 1993, Noctor et al. 2014). ^1^O_2_ is a highly reactive form of oxygen that causes oxidative damage to key cellular components, including lipids in the thylakoid membranes, proteins and pigments in the photosynthetic complexes, and nucleic acids, thereby compromising cellular functions (Triantaphylidès and Havaux 2009, Lee and Kim 2024).

Carotenoids, which are located in the photosystems near the chlorophyll molecules, constitute the first line of defense against ^1^O_2_ toxicity in plant leaves (Caferri et al. 2022, Sun et al. 2023). ^1^O_2_ quenching by carotenoids mainly proceeds via a physical mechanism that results in the dissipation of thermal energy (Edge and Truscott 2018). However, since carotenoids comprise multiple isoprene units with alternating conjugated double bonds, they are susceptible to oxidation by ^1^O_2_ or enzymatic action (Nisar et al. 2015). Consequently, carotenoids can also quench ^1^O_2_ by a chemical quenching mechanism that involves direct oxidation of the carotenoid molecule by ^1^O_2_ (Stratton et al. 1993, Ramel et al. 2012a). This phenomenon leads to a variety of oxidized derivatives called apocarotenoids. A crucial site of ^1^O_2_ production is the β-carotene-containing PSII reaction center (Krieger-Liszkay 2005, Pospisil 2016), which is also a major generator of oxidized β-carotene metabolites such as β-cyclocitral (β-CC) (D’Alessandro and Havaux, 2019). The basal level of this apocarotenoid in leaves (approximately 50 ng g^−1^ fresh weight) can triple under conditions of high light stress or drought (Ramel et al., 2012b, D’Alessandro et al. 2019).

β-CC is a volatile compound that acts as a signal molecule in plants (Havaux 2020, Deshpande et al. 2021). This compound triggers changes in the expression of ^1^O_2_-responsive genes, facilitating acclimatization to ^1^O_2_ and photooxidative stress (Ramel et al. 2012b). β-CC is among several signaling metabolites recently linked to chloroplast-to-nucleus retrograde signaling (e.g. Chan et al. 2016). β-CC is partially converted to β-cyclocitric acid (β-CCA) *in planta* (D’Alessandro et al. 2019), constituting one of the initial steps in β-CC-dependent signaling. In fact, under drought conditions, there is a 15- fold increase in β-CCA concentration (D’Alessandro et al. 2019). The expression of β-CC- and ^1^O_2_ - responsive genes was induced when plants were exposed to exogenous β-CCA (D’Alessandro et al. 2019). Gene upregulations induced by β-CC and β-CCA were dominated by the induction of the xenobiotic detoxification pathway that eliminates toxic compounds produced during oxidative stress (Braat et al. 2023, D’Alessandro et al. 2018). As a result, β-CC and β-CCA markedly enhance plant resistance to abiotic stress conditions (Braat et al. 2023, D’Alessandro et al. 2018 and 2019, Zhu et al. 2023). However, the β-CC signaling mechanism appears to be more complex, possibly involving a multibranched signaling cascade. Indeed, the protection provided by β-CC was found to be dependent on the zinc finger protein MBS1 (Shumbe et al. 2017, D’Alessandro et al. 2018) whereas the effects of β-CCA were insensitive to the lack of MBS1 (D’Alessandro et al. 2019), indicating the existence of a signaling pathway specific to β-CC.

We have undertaken the present study to characterize the signaling pathway that involves both β-CC and β-CCA. The transcriptomic profiles of Arabidopsis leaves after exposure to β-CC or β- CCA over different exposure times were compared to identify differentially expressed genes (DEGs) that are common to these conditions. Of these, the most-induced DEG under all conditions was a gene that belongs to the cytochrome P450 family (CYP). We show that this *CYP* gene is a crucial component of the stress resistance induced by β-CC and β-CCA and that it acts in particular through a modulation of leaf photosynthetic capacity.

## Results

### Transcriptomic analysis of the effects of β-CC and β-CCA on Arabidopsis leaves

β-CC and its oxidized derivative β-CCA are both signaling molecules that induce changes in nuclear gene expression leading to plant stress tolerance (Ramel et al. 2012b, D’Alessandro et al. 2019, Havaux 2020). To investigate the signaling pathway involving both compounds, we performed a RNAseq analysis of gene expressions in leaves of Arabidopsis plants treated with β-CC or β-CCA over 2 exposure times (4 h and 9 h). Compared to watering with a β-CCA solution, the treatment of Arabidopsis plants with volatile β-CC affected the expression of more genes (Fig. 1A). This difference is possibly due, at least in part, to different modes of application of β-CC and β-CCA (volatile β-CC in the gas phase around the plants placed in a closed box vs. watering of the soil with β-CCA) which lead to different apocarotenoid concentrations in the leaves (D’Alessandro et al. 2019). Exposure to β-CC for 4 and 9 h changed the expression of 1823 and 5298 genes, respectively. 4-h exposure to β-CCA impacted the expression of 689 genes. Rather surprisingly, the number of DEGs dropped to 59 after 9 h of β-CCA treatment, suggesting a more transient effect of β-CCA on gene expression relative to β-CC. This could be related to plant metabolization of exogenously applied β-CCA (D’Alessandro et al. 2019).

**Fig. 1.**
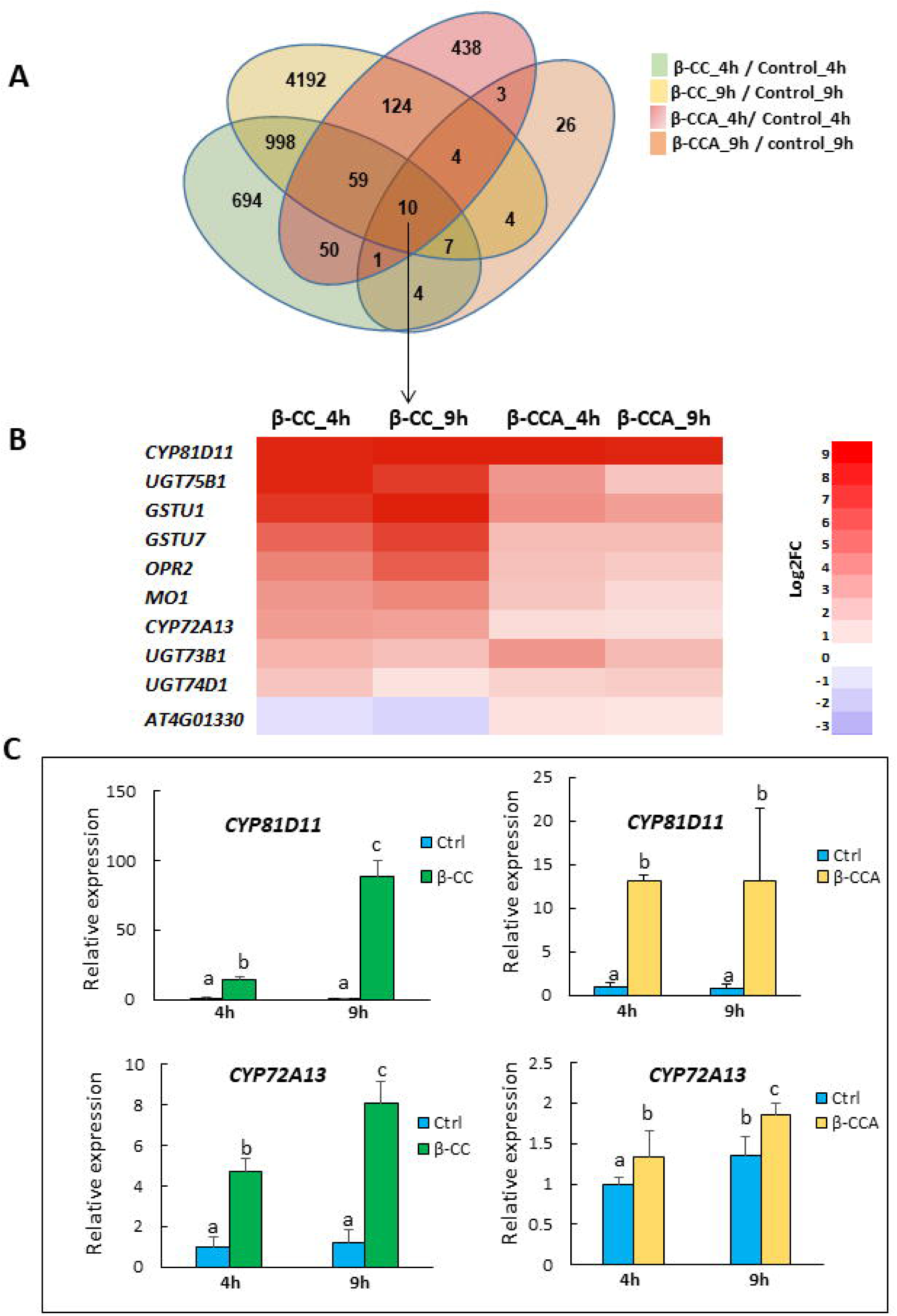
RNAseq and qRT-PCR analyses of gene expressions in leaves of Arabidopsis plants treated with β-CC or β-CCA. A) RNAseq: Venn diagram of genes differentially expressed (DEGs) in Arabidopsis leaves exposed to β-CC or β-CCA for 4 h or 9 h compared to their respective control leaves (control 4 h and 9 h). Data are average values of 3 independent samples. B) RNAseq: Heat map of the expression of the 10 DEGs common to all conditions. C) qRT-PCR analysis of the expressions of the CYP81D11 and CYP72A13 genes in leaves treated for 4 h or 9 h with β-CC or β-CCA. Data are expressed relative to control 4 h and are average values of at least 6 repetitions + SD. Different letters indicate significant difference at P< 0.05 (Anova – Duncan’s test).

Only 10 genes were common to all conditions (Fig. 1B). Most of those genes encode proteins related to cellular detoxification processes such as glutathione transferase (2 *GST* genes), glucosyl transferase (2 *UGT* genes), cytochrome P450 (2 *CYP* genes) and OPR2. The latter protein, which belongs to the family of OPDA (12-oxo-phytodienoic acid) reductases but is not involved in jasmonate synthesis, has been reported to play a physiological role in detoxification of xenobiotics such as trinitrotoluene (Beynon et al. 2009). The two other genes encode a monooxygenase (*MO1*) and a putative kinase (*AT4G01330*) of unknown function. The latter gene behaved differently from the other genes: it was downregulated by β-CC and slightly upregulated by β-CCA, contrary to all the other genes which were upregulated by both β-CC and β-CCA. Because of its peculiar response to β- CC, this gene was not considered further in this study.

*CYP81D11* (*AT3G28740*) was the most induced gene under all conditions (log_2_ (fold changes FC) > 8, Fig. 1B). The other genes were induced by β-CC to variable extent (but less than *CYP81D11*) and were induced only weakly by β-CCA. This was confirmed by qRT-PCR analyses of the two CYP genes (Fig. 1C). *CYP81D11* was markedly induced by β-CC (up to x100 after 9 h) and β-CCA (around x12). In contrast, *CYP72A13* (*AT3G14660*) was very weakly induced by β-CCA (around x1.5) and moderately by β-CC (x8) compared to *CYP81D11*. qRT-PCR analyses also confirmed that the other genes listed in Fig. 1B are responsive to β-CC and β-CCA, with lower induction levels compared to CYP81D11 (Figs. S1 and S2).

### The genes induced by both β-CC and β-CCA are controlled by TGAII transcription factors

The xenobiotic detoxification pathway is known to be controlled by Class II TGA transcription factors (TGAII) interacting with the SCL14 (SCARECROW LIKE 14) transcription regulator (Mueller et al. 2008, Fode et al. 2008). Interestingly, all the 10 genes identified in Fig. 1B contain the TGACG binding motif (Table S1), suggesting that their induction by β-CC and β-CCA is under the control of TGAII transcription factors. This is confirmed by a comparative qRT-PCR analysis of WT Arabidopsis and the triple mutant *tga256* (Fig. 2, Figs. S1 and S2). Lack of TGAII-2, TGAII-5 and TGAII-6 in tga256 drastically inhibited the expression of the *CYP81D11* gene before and after the β-CC and β-CCA treatments (Fig. 2A and 2B). *CYP72A13* expression was also perturbed in *tga256*, but the effects were less pronounced (Fig. 2C and Fig. 2D). In particular, under control conditions, the expression of the *CYP72A13* gene appeared to be weakly dependent on TGAII. The inhibition of the β-CC and β-CCA effects on the expression of this gene in *tga256* was also less marked compared to the CYP81D11 expression changes. The expression of most genes identified in Fig. 1B was also deregulated by the *tga256* mutations in a way similar to *CYP72A13*: no or weak effect in control conditions and inhibition of the β-CC- and β-CCA-induced upregulations (Fig. S1 and Fig. S2). However, contrary to the other genes, the expression of *GSTU1, GSTU7* and *UGT73B1* under control conditions was substantially reduced in the tga256 mutant compared to WT.

**Fig. 2.**
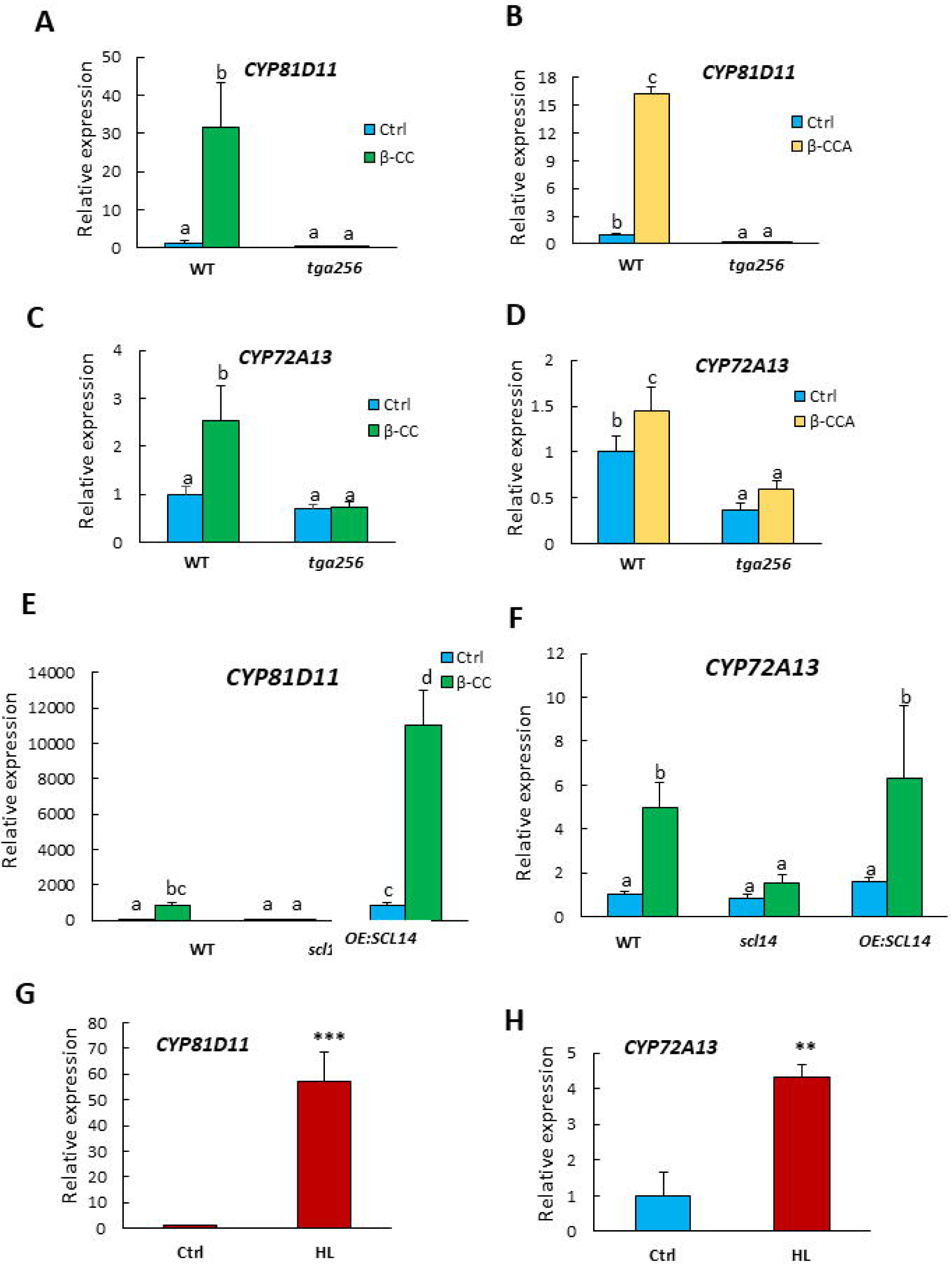
The expression of the CYP genes induced by both β-CC and β-CCA is controlled by TGAII/SCL14 transcription factors/regulator and is induced by high light. A-D) qRT-PCR analysis of expression of the CYP81D11 and CYP72A13 genes in Arabidopsis WT and tga256 triple mutant under control conditions and after treatment with β-CC or β-CCA. E-F) qRT-PCR analysis of the expression of CYP81D11 and CYP72A13 genes in WT, scl14 knockout mutant and SCL14-overexpressing line (OE:SCL14) under control conditions and after treatments with β-CC. G-H) Expression levels of the CYP81D11 and CYP72A13 genes in response to high light (HL, 1500 µmol photons m s for 3 h). Data are expressed relative to control WT (Ctrl) and are average values of at least 6 repetitions + SD. *** and **, significant difference at P< 0.001 and P< 0.01, respectively (Student’s t-test). Different letters indicate significant difference at P< 0.05 (Anova – Duncan’s test).

We also examined the role of the SCL14 regulator in the response of CYP81D11 and CYP72A13 to β-CC using the *scl14* knockout mutant previously described (Fode et al. 2008, D’Alessandro et al. 2018). Similarly to the tga256 triple mutant, the scl14 mutant showed virtually no induction of *CYP81D11* expression by β-CC (Fig. 2E). On the opposite, in the *SCL14*-overexpressing *OE:SCL14* line, there was a spectacular increase in *CYP81D11* expression both under control conditions and after the β-CC treatment. These effects were much less marked for CYP72A13 (Fig. 2F). The expression of the latter gene in *scl14* remained unchanged under control conditions and was inhibited, but partially only, after the β-CC treatment. Moreover, overexpressing SCL14 had little effect on CYP72A13 expression. The other genes in the list of Fig. 1B were also dependent on SCL14 (Fig. S3). The expression profile of *UGT74D1, UGT73B1, GSTU7* and *OPR2* in the scl14 mutant resembled that of CYP72A13 whereas UGT75B1, GSTU1 and MO1 behaved almost like CYP81D11.

To sum up, the *CYP81D11* gene is strongly induced by β-CC and β-CCA and is under the strict control of the transcription-regulating TGAII-SCL14 complex. *CYP72A13* and the other genes of Fig. 1B are less responsive to β-CC and β-CCA with a weaker dependence on TGAII-SCL14. Because of the remarkable responses of CYP81D11 to β-CC and β-CCA, the present study will focus mainly on this gene.

### The CYP81D11 overexpressor is highly tolerant to photooxidative stress

β-CC and β-CCA accumulate in Arabidopsis leaves under high light stress and drought stress (Ramel et al. 2012b, D’Alessandro et al. 2019). The expression of CYP81D11 was measured by qRT-PCR in response to both conditions. *CYP81D11* was strongly induced (X60) after 3 h in high light (Fig. 2G). This gene was also induced by drought (no watering for 7 d), but the induction was much less pronounced compared to the effect of high light (Fig. S4A). Similarly to what we observed in response to β-CC and β-CCA, the high light-induced changes in CYP72A13 expression occurred in an attenuated form compared to CYP81D11 (Fig. 2H). *CYP72A13* was also inducible by drought conditions, and the amplitude of this effect was low and comparable to the *CYP81D11* expression changes under this condition (Fig. S4B).

Block of the xenobiotic detoxification pathway in the tga256 mutant is associated with an increased sensitivity to photooxidative stress (D’Alessandro et al. 2018). It is known that the xenobiotic detoxification pathway targets toxic molecules generated during oxidative stress such as lipid-derived reactive carbonyl species (RCS) (Mano et al. 2019). This is confirmed in Fig. 3: High light stress led to an accumulation of lipid peroxides (Fig.3B) and lipid RCS (Fig. 3C) compared to WT, indicating enhanced oxidative damage to lipids and subsequent degradation of the lipid peroxides. As expected (D’Alessandro et al. 2018), β-CC markedly reduced lipid damage in high light-treated WT plants, but this protective effect of β-CC was absent in tga256 (Fig. 3A-C).

**Fig. 3.**
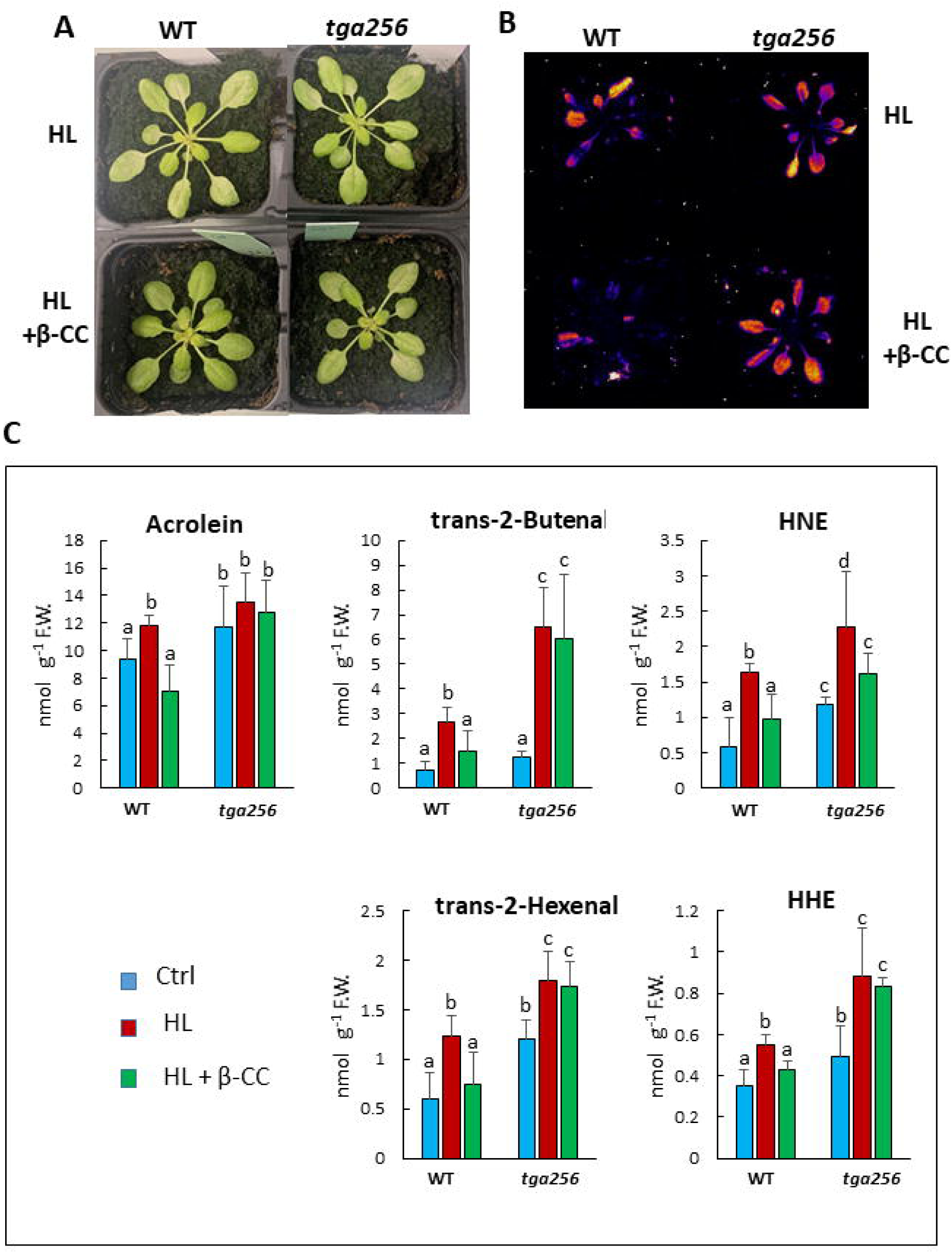
Relative sensitivity of Arabidopsis WT and tga256 triple mutant plants to phototoxidative stress in response to β-CC. Plants were exposed to high light (HL, 1500 µmol photons m^-2^ s^-1^) at low temperature (9°C) for 2 d. Some plants were pretreated with volatile β-CC for 4 h. A) Picture of the plants after high light stress. B) Autoluminescence images of lipid peroxidation in whole plants after high light stress. C) RCS levels in leaves before and after high light stress. Data are mean values of 4 to 8 repetitions + SD. Different letters indicate significant differences at P< 0.05 (Anova – Duncan’s test).

The same stress treatments were applied to a CYP81D11-overexpressing Arabidopsis line (*OE:CYP81D11*, line 12-2.1) (Fig. 4). Interestingly, *OE:CYP81D11* plants were found to be very resistant to high light stress with a marked reduction in lipid peroxidation and RCS levels compared to WT (Fig. 4A-D). This high phototolerance was confirmed in a number of other overexpressing lines (Fig. S5). Thus, CYP81D11 appears to be an important component of plant tolerance to photooxidative stress. Accordingly, a cyp81d11 knockout Arabidopsis mutant exhibited a phenotype opposite to that of the overexpressors (Figs. 4E-G): *cyp81d11* plants were more photosensitive than WT plants, with more lipid peroxidative damage after a high light stress treatment (Figs. 4F-G). For comparison purposes, we also examined the responses of *OE:CYP72A13* lines to high light stress. Contrary to *CYP81D11* overexpression, overexpressing *CYP72A13* did not enhance high light tolerance (Fig. S6).

**Fig. 4.**
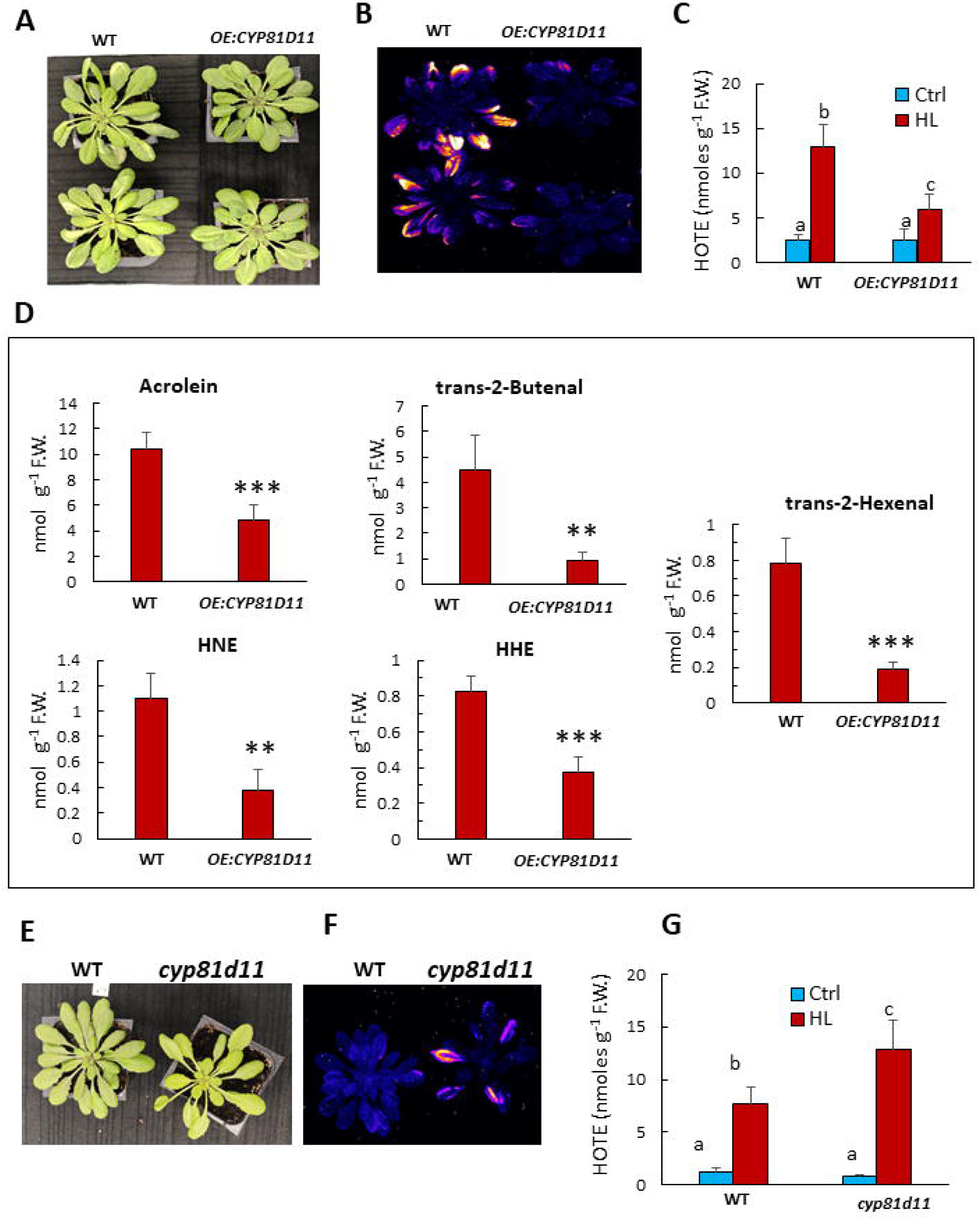
Relative sensitivity of Arabidopsis WT, **OE:CYP81D11** and *cyp81d11* plants to photooxidative stress. Plants were exposed to high light (HL, 1500 µmol photons m^-2^ s^-1^) at low temperature (9°C) for 2 d. A) Picture of the WT and *OE:CYP81D11* after high light stress. B) Autoluminescence images of lipid peroxidation in whole plants after high light stress. C) HOTE levels in WT and *OE:CYP81D11* leaves before and after high light stress. Data are mean values of at least 3 repetitions + SD. D) RCS levels in WT and **OE:CYP81D11** leaves after high light stress. Data are mean values of 4 to 8 repetitions + SD. E) Picture of the WT and cyp81d11 after high light stress. F) Autoluminescence images of lipid peroxidation in whole WT and cyp81d11 plants after high light stress. G) HOTE levels in WT and cyp81d11 leaves before and after high light stress. Data are mean values of at least 3 repetitions + SD. *** and **, significant difference at P< 0.001 and P< 0.01, respectively (Student’s t-test). Different letters indicate significant difference at P< 0.05 (Anova – Duncan’s test).

We were interested to check the effects of drought, another pro-oxidative condition that can enhance ROS photoproduction including singlet oxygen (Smirnoff 1993, Noctor et al. 2014, Chen and Fluhr 2018). *OE:CYP81D11* lines were noticeably more tolerant to drought stress than WT (Fig. S4C and D). On the opposite, *cyp81d11* mutant plants were more sensitive to drought stress than WT plants, with a more reduced RWC (Relative Water Content) after 7-d of water deprivation (Fig. S4E- F). However, since drought had relatively little effects on CYP81D11 expression (Fig. S4A), the CYP81D11 levels reached in drought-stressed WT plants might be very low, possibly limiting its role in the modulation of plant drought tolerance under natural conditions.

*OE:CYP72A13* lines exhibited an increased tolerance to water stress, with a better preservation of leaf water content, compared to WT (Fig. S7). Nevertheless, *OE:CYP72A13* plants did not seem to reach the drought tolerance levels of *OE:CYP81D11* plants (Fig. S7 vs. Fig. S4).

### CYP81D11 overexpression does not lead to a stimulation of the detoxification pathway

Cytochrome P450 are hemoproteins that function mostly as monooxygenases, being involved in various NADPH- and O_2_-dependent hydroxylation reactions that produce a variety of plant compounds (Bak et al. 2011). The reaction catalyzed by CYP81D11 is not known. Plants treated with volatile β-CC accumulate β-CC in their leaves which is partially oxidized *in vivo* to β-CCA (D’Alessandro et al. 2019). We measured the concentrations of β-CC and β-CCA in β-CC-treated Arabidopsis plants (Fig. S8). No significant difference was observed between WT and **OE:CYP81D11** which accumulate the same amounts of β-CC and β-CCA. It can therefore be concluded that CYP81D11 is not involved in the biosynthesis or metabolization of the two apocarotenoids.

Based on its protective effects against abiotic stresses and RCS accumulation, we can imagine that this cytochrome somehow modulates the whole detoxification pathway through a signaling effect. We checked this possibility by measuring the expression of a number of genes known to be involved in the β-CC pathway including several detoxification genes (*SDR1, AER, MAPKKK18, TOLL LIKE INTERLEUKIN, ANAC102, GSTU5*) using qRT-PCR (Fig. 5). None of the examined genes were more induced by β-CC in the CYP81D11 overexpressor compared to WT. Rather surprisingly, the expression of several genes (*AER, GSTU5, MAPKKK18*) was even much less induced in **OE:CYP81D11** relative to WT. One can conclude that the increase in stress tolerance associated by high levels of CYP81D11 does not rely on a general stimulation of the xenobiotic detoxification pathway. This conclusion is also supported by RNAseq analyses of **OE:CYP81D11** (see below).

**Fig. 5.**
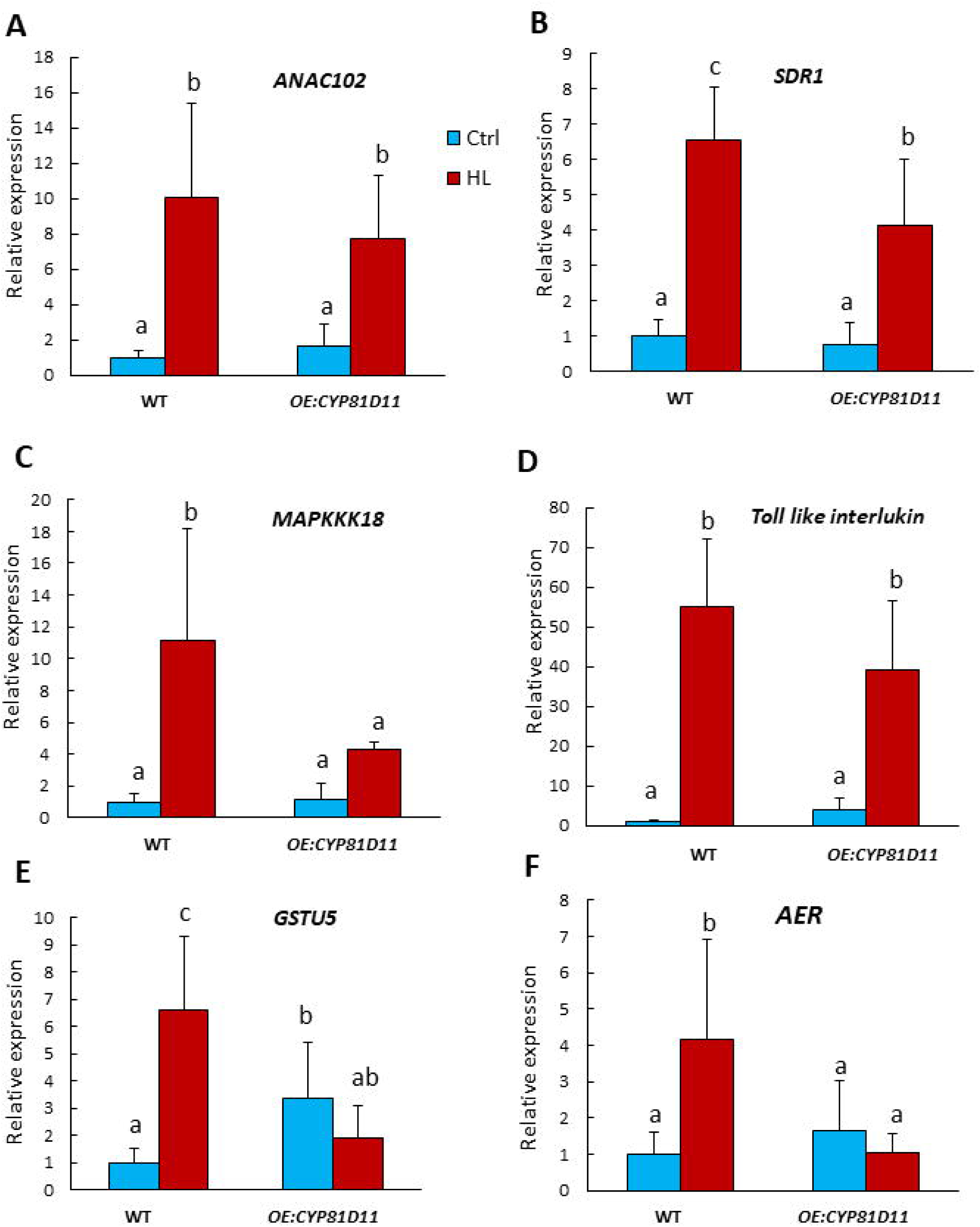
Effects of CYP81D11 overexpression on the expression of β-CC-responsive genes. qRT-PCR analyses were performed on Arabidopsis WT and **OE:CYP81D11** leaves before and after high light stress (HL, 1500 µmol photons m^-2^ s^-1^ for 3 h). Data are expressed relative to control (ctrl) WT and are average values of at least 6 repetitions + SD. Different letters indicate significant difference at P< 0.05 (Anova – Duncan’s test).

### Transcriptomic analysis of CYP81D11 overexpression points to changes in photosynthesis-related genes

To have a more complete picture of the transcriptomic consequences of *CYP81D11* overexpression, a RNAseq analysis was performed to compare gene expressions in the overexpressor vs. WT (grown under control conditions). Around 1800 genes were upregulated in *OE:CYP81D11* while 1900 genes were downregulated relative to WT (Fig. 6A). Fig. 6B shows the functional categories of the most affected genes. DEGs essentially belong to two main categories: photosynthesis (photosynthesis, light harvesting, photosystem II assembly, thylakoid membrane organization, photosynthetic electron transport,…) and defense against stresses (oxidative stress, cold, hypoxia,…). No categories related to cellular detoxification or metabolism and transport of toxic compounds are present in the list of functions in Fig. 6B, thus confirming the qRT-PCR analyses of Fig. 5 which did not show stimulation of detoxification-related genes.

**Fig. 6.**
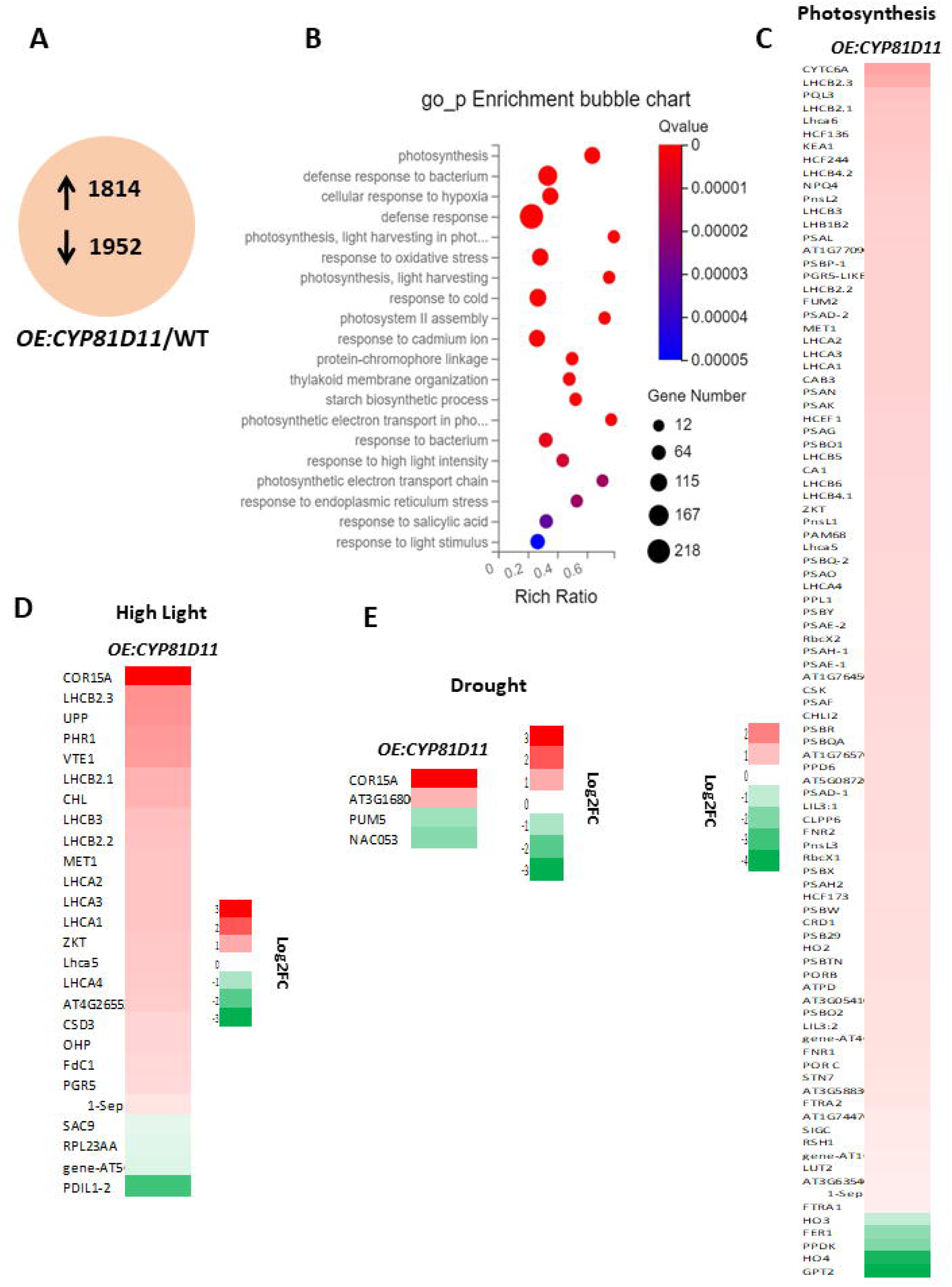
RNAseq transcriptomic analysis of **OE:CYP81D11** compared to WT. Plants were grown under normal control conditions. A) Number of DEGs (DESeq, P-adjusted <0.05, |log_2_ (fold change)| ≥0) in leaves of **OE:CYP81D11** versus WT. Upwards and downwards arrows indicate up-regulation and down-regulation, respectively. Data are average values of 3 experiments. B) Gene ontology biological process (GOPB) enrichment bubble plots. Functional categories of the most affected genes by CYP81D11 overexpression are shown. Bubble color and size indicate the Q value and gene number enriched in the biological process. C-E) Heat maps of DEGs in **OE:CYP81D11** leaves compared to WT plants. Genes are related to C) photosynthesis, D) response to high light stress and E) response to drought. P < 0.05. Log_2_ (fold change) ≥0.

In general, the relative changes in gene expression in **OE:CYP81D11** are rather moderate. The heat maps of Figs. 6 C-E detail the expression changes of genes related to photosynthesis, to high light stress and to drought. Strikingly, COR15A (AT2G42540) stands out from the rest of the genes in the ‘high light stress’ category, showing a marked increase in expression (Figs. 6D). The number of DEGs in the ‘response to drought’ section was very limited, but again induction of COR15A was a striking feature of this functional category (Fig. 6E). These are interesting observations because COR15A is a stress-inducible protein that can protect chloroplasts (Artus et al. 1996, Thalhammer et al. 2014). This category includes also the gene encoding the plastid lipocalin (CHL, also named LCNP, AT3G47860) which protects thylakoid membranes lipids against oxidation under various stresses including high light stress (Levesque-Tremblay et al. 2009) (Fig. 6D). Concerning photosynthesis, many genes coding for proteins of the PSII and PSI reaction centers (psb and psa genes, respectively) and antennae (*Lhcb* and *Lhca* genes, respectively) were upregulated. The levels of this upregulation were, however, rather modest (log_2_ FC <2) (Fig. 6C).

### CYP81D11 enhances leaf photosynthetic capacity and plant growth in high light

Since high expression levels of CYP81D11 appear to affect genes related to the chloroplast (COR15A, CHL, photosynthesis-related genes), we measured the efficiency of photosynthetic electron transport in **OE:CYP81D11** and WT leaves (grown under control conditions) by *in vivo* chlorophyll fluorometry. The plot of Φ_PSII_ vs. PFD (Fig. 7A) shows that photosynthetic electron transport was much less rapidly saturated by light in **OE:CYP81D11** leaves relative to WT leaves. The quantum yield of PSII photochemistry was almost doubled in the overexpressor at high PFDs above 1500 µmol photons m^-2^ s^-1^. In low light (e.g. at the growth PFD), the two genotypes did not significantly differ. Acclimation of photosynthesis to high light conditions may have various causes including a decrease in light absorption. We can exclude a stimulation of energy dissipation in the photosystems since NPQ was not enhanced in **OE:CYP81D11** compared to WT (Fig. 7B).

**Fig. 7.**
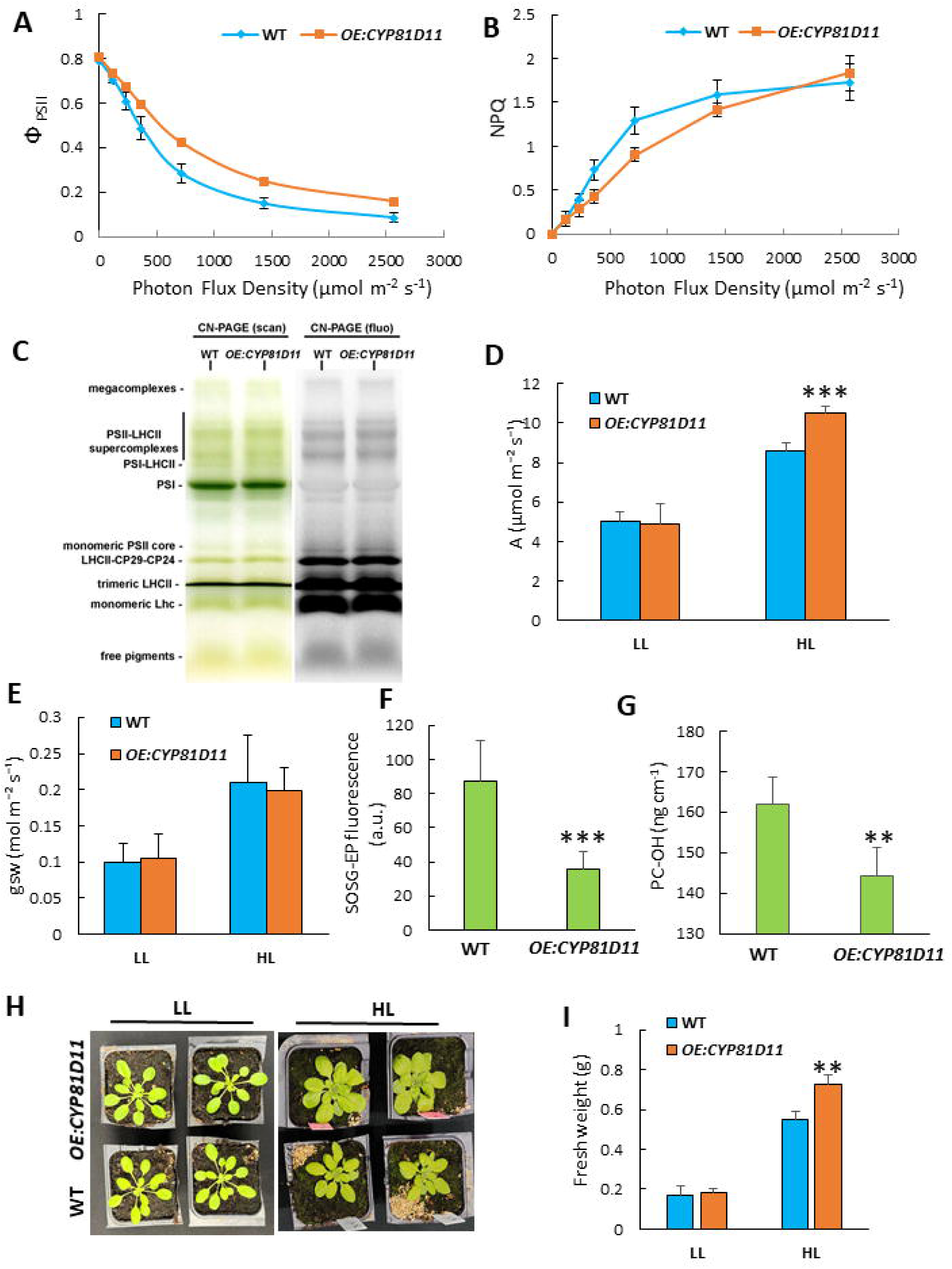
Photosynthesis and ^1^O_2_ photoproduction in Arabidopsis WT and **OE:CYP81D11** plants. A-E) Plants were grown under control conditions. A) Efficiency of photosynthetic electron transport (Φ_PSII_) at different PFDs of white light. B) NPQ at different PFDs. A and B) Data are average values of 4 repetitions ± SD. C) CN-PAGE separation of photosynthetic complexes. Left: native gel; Right: fluorescence image of the gel (excitation, 470 nm; emission, 690-720 nm), allowing a better visualization of the highly fluorescent PSII and LHC antenna complexes. D) CO_2_ fixation rate A by WT and **OE:CYP81D11** leaves in low light (LL, 130 µmol photons m^-2^ s^-1^) and in high light (HL, 1500 µmol m^-2^ s^-1^). E) Stomatal conductance gsw of WT and *OE:CYP81D11* leaves in low light and in high light. D and E) data are average values of 3 to 6 repetitions + SD. F-G) ^1^O_2_ production in high light (20 min at PFD 1500 µmol m^-2^ s^-1^) as measured by F) the fluorescence intensity of SOSG-EP and G) the levels of hydroxyl-plastochromanol (PC-OH). Data are mean values of 4 repetitions + SD. H) Pictures of WT and *OE:CYP81D11* plants grown for 21 d in low light or in high light (130 or 1100 µmol photons m^-2^ s^-1^, respectively). I) Plant biomass after 21-d growth in low light or in high light (130 or 1100 µmol photons m^-2^ s^-1^, respectively). Data are mean values of 4 repetitions. *** and **, significant difference at P< 0.001 and P< 0.01, respectively (Student’s t-test).

We measured the concentrations of chlorophyll a and chlorophyll b in WT and *OE:CYP81D11* leaves (Fig. S9). Since chlorophyll b is exclusively present in the light-harvesting chlorophyll antennae of the photosystems, a decrease in the antenna size will be reflected by a higher chlorophyll *a/b* ratio. In fact, we observed a parallel increase in both chlorophylls in *OE:CYP81D11* relative to WT, with a very small decrease in the chlorophyll a/b ratio (from 4.1 to 3.9). It should be remembered that a marked increase in chlorophyll *a/b* ratio is disadventageous in low light since it corresponds to a decreased light absorption capacity (less chlorophyll b-containing light harvesting antennae). The similar Φ_PSII_ values in WT and *OE:CYP81D11* leaves in low light is thus consistent with limited effects of CYP81D11 overexpression on the light-harvesting antenna size. This conclusion was confirmed by an analysis of the composition of photosynthetic complexes in WT and *OE:CYP81D11* thylakoids (Fig. 7C). After solubilization from thylakoid membranes, photosynthetic complexes were separated by Clear Native Polyacrylamide gel electrophoresis (CN-PAGE). The profiles obtained for the two genotypes were very similar. In particular, the abundance of the PSII light-harvesting antennae was not changed by the overexpression of *CYP81D11*. We can conclude that changes in the photosynthetic complex composition of *OE:CYP81D11* thylakoids, if any, must be very subtle.

Increased efficiency of photosynthetic linear electron flow could also result from the removal of a limitation at the PSI acceptor side, e.g. due to a stimulation of the dark reactions of CO_2_ assimilation leading to a faster consumption of photosynthesized ATP and NADPH. CO_2_ fixation was measured using an infrared gas analyzer (IRGA). The rate of leaf photosynthesis (A) in low light (130 µmol photons m^-2^ s^-1^) did not differ between WT and *OE:CYP81D11* (Fig. 7D). In contrast, when PFD was increased to 1500 µmol photons m^-2^ s^-1^, the rate of CO_2_ fixation was noticeably higher in *OE:CYP81D11* leaves relative to WT leaves. Stomatal conductance (gsw) was observed to increase when leaves were exposed to high light (Fig. 7E), and this effect occurs in both WT and *OE:CYP81D11*. We did not find any significant difference in gsw between the two genotypes both in low light and in high light.

A higher photosynthetic efficiency in high light could reduce ROS production in the chloroplasts. We measured the photoproduction of ^1^O_2_ in WT and *OE:CYP81D11* using the SOSG fluorescence probe (Flors et al. 2006, Prasad et al. 2018). Fig. 7F shows that the fluorescence of SOSG-endoperoxide (SOSG-EP), measured with a fiberoptics-equipped spectrofluorometer after 20- min illumination, noticeably decreased in *OE:CYP81D11* leaves compared to WT leaves. We also quantified the leaf content in plastochromanol-OH (PC-OH), a specific marker of ^1^O_2_ oxidation (Szymanska et al. 2014, Kruk and Szymanska 2021), by HPLC coupled with fluorescence detection, and we found higher levels in WT leaves (Fig. 7G). These results indicate that *OE:CYP81D11* produced less ^1^O_2_ in high light than WT.

To determine the effects of the photosynthetic changes associated with *CYP81D11* overexpression on plant growth, WT and *OE:CYP81D11* plants were grown for 21 d in low light (130 µmol photons m^-2^ s^-1^) or in strong light (1100 µmol m^-2^ s^-1^) at 21°C air temperature (Fig. 7H-I). No significant difference in plant biomass was observed between the two genotypes in low light. In contrast, at high PFD, the size (Fig. 7H) and biomass (Fig. 7I) of *OE:CYP81D11* plants was noticeably higher (> +30%) than that of WT plants. Thus, high CYP81D11 levels are advantageous for plant biomass productivity, especially in high light environments.

## Discussion

The CYP81D11 gene has been previously reported to be inducible by some oxylipins, such as cis-jasmone, which can be produced by insect-induced leaf wounding (Bruce et al. 2008, Stotz et al. 2013, Walper et al. 2016). In addition, there are some reports correlating the expression levels of this gene and plant resistance to herbivorous insects (Bruce et al. 2008, Matthes et al. 2010). The present study shows that CYP81D11 is also induced by abiotic stress conditions, such as high light and drought, and that this induction can be mediated by signaling apocarotenoids, such as β-CC and β-CCA, generated in plant leaves under those environmental constraints. The common feature of the molecular inducers of the CYP81D11 gene under biotic and abiotic stresses is their reactivity associated with their electrophilic properties (Havaux 2020, Farmer and Mueller 2013). For instance, β-CC and cis-jasmone contain an α-β unsaturated carbonyl group which is known to confer a high electrophilicity to the molecules allowing them to bind to nucleophilic atoms of macromolecules (Farmer and Mueller 2013). However, the effect of β-CC on gene expression cannot be merely considered as a general response to reactive electrophile species since another electrophilic β- carotene derivative, β-ionone, showed a low efficiency in inducing β-CC-responsive genes (Ramel et al. 2012b), pointing to specific responses and to a possible role of some structural factors in the signaling activity of apocarotenoids.

Reactive electrophile species are known to reprogram gene expression through a TGAII transcription module, changing the expression of a number of genes, especially genes encoding detoxification functions and cell cycle regulators (Farmer and Mueller 2013). Accordingly, β-CC-induced expression of CYP81D11 was found to be entirely dependent on TGAII and its regulator SCL14. Endogenous toxic compounds targeted by the TGAII-controlled detoxification pathway include products derived from lipid peroxidation (Farmer and Mueller 2013). Consistently, the tga256 triple mutant accumulated much more lipid RCS than WT after photooxidative stress. On the opposite, the CYP81D11 overexpressor contained less RCS than WT after high light stress. However, we cannot directly interpret the latter finding in terms of increased detoxification capacities since high light-induced lipid peroxidation was much lower in *OE:CYP81D11* compared to WT, likely producing less secondary reactive by-products.

A striking result was that high expression levels of CYP81D11 led to a marked enhancement of Arabidopsis tolerance to photooxidative stress and drought stress. In fact, induction of CYP81D11 expression was sufficient to allow plants to reach stress tolerance levels comparable to the protection provided by β-CC and β-CCA (D’Alessandro et al. 2018, 2019), suggesting that CYP81D11 is a crucial component of apocarotenoid-induced stress tolerance. On the opposite, an Arabidopsis CYP81D11-deficient mutant was less tolerant to high light relative to WT, in line with this interpretation.

Cytochromes P450 form a very large family of proteins implied in a variety of metabolic processes (Hansen et al. 2021, Pandian et al. 2020, Werck-Reichhardt and Feyereisen 2000). The most common reaction catalyzed by cytochromes P450 is a monooxygenase reaction. They catalyze NADPH-and/or O_2_-dependent hydroxylation reactions in primary and secondary metabolism, contributing to the diversity of plant metabolites, to xenobiotic metabolism and to biosynthesis of defense compounds. The reaction catalyzed by CYP81D11 is unfortunately unknown, as is that of most CYP81 proteins. The Arabidopsis genome encodes 17 genes of the *CYP81* family, including 9 CYP81Ds, of which very few have been subjected to functional analysis (Bak et al. 2011, Wang et al. 2020). We can exclude the participation of CYP81D11 in the oxidation of β-CC to β-CCA in planta since no difference in the concentration of those apocarotenoids was observed between WT and *OE:CYP81D11* leaves. It is also unlikely that the high stress tolerance of *OE:CYP81D11* plants is due to the direct involvement of the cytochrome P450 in a crucial, yet uncharacterized, detoxification reaction that would eliminate the bulk of reactive and toxic molecules generated in the stressed leaves. Indeed, the chemical nature of the toxic molecules produced during lipid peroxidation is known to be very diverse requiring a large panoply of detoxifying enzymes catalyzing specific reactions (Sousa et al. 2017, Biswas and Mano 2021, Moldogazievza et al. 2023, Mano et al. 2019). It is very unlikely that the enzymatic activity of CYP81D11 alone would be able to carry out all these detoxification reactions.

CYP81D11 does not appear either to be a key element in the apocarotenoid signal transduction. Indeed, overexpressing *CYP81D11* had no stimulatory effect on the expression of β-CC-and β-CCA-responsive genes. We can therefore suggest that CYP81D11 accumulation somehow enhances some additional molecular mechanisms that contribute to plant stress tolerance, supplementing cellular detoxification and reducing oxidative stress and lipid peroxidation. A comparative transcriptomic analysis of WT and *OE:CYP81D11* confirmed that the predominant effect of overexpressing *CYP81D11* was not on detoxification processes or cell cycle regulations. High CYP81D11 expression level was mainly associated with changes in photosynthesis-and high light stress-related genes. Remarkably, the efficiency of photosynthetic electron transport and the capacity of photosynthetic CO_2_ fixation were enhanced in *OE:CYP81D11* leaves compared to WT leaves, and this effect was especially visible in high light. Because the number of photosynthesis-related genes upregulated in the CYP81D11 overexpressor was high and the extent of the upregulation was rather uniform and moderate, it is difficult at this stage of our study to determine the specific modifications that led to this photosynthetic improvement. However, based on our CN-PAGE analysis of thylakoid protein complexes, we can exclude remodeling of the photosynthetic complex composition in *OE:CYP81D11* thylakoids.

Photosynthesis was stimulated in *OE:CYP81D11* leaves in high light, not in low light, that is under conditions were light is not a limiting factor. Improvement of the electron transport efficiency in light-saturating conditions could be due to an electron sink that removes photochemical limitations, e.g. at the PSI acceptor side. Interestingly, expression of *CYP1A1* in cyanobacteria was previously reported to enhance Φ_PSII_ specifically in high light and this phenomenon was interpreted as a change in source/sink imbalance (Santos-Merino et al. 2021). One possibility could be an effect of CYP81D11 on the biochemical ‘dark’ phase of photosynthesis, favoring consumption of ATP and the reducing power. CO_2_ supply to the chloroplasts does not seem to be the target of CYP81D11 since there was no difference in stomatal conductance between WT and *OE:CYP81D11* leaves in either low light and high light. Therefore, one can imagine a role for CYP81D11, either direct or indirect, in the regulation of the biochemical reactions of photosynthetic CO_2_ assimilation. Alternatively, high CYP81D11 levels could generate a reductant-consuming metabolic pathway that remains to be established. However, a major rerouting of electrons to an alternative, non-photosynthetic metabolic pathway does not seem to be in line with the increase in CO_2_ photoassimilation activity observed in *OE:CYP81D11*. It is clear that an important challenge for future work will be to investigate these possibilities and to determine how CYP81D11 can modulate the rate of photosynthetic CO_2_ fixation. Irrespective of the mode of action of CYP81D11 on photosynthesis, an increase in photosynthesis efficiency in high light can reduce ROS photoproduction (Lee and Kim 2024) and can therefore enhance leaf tolerance to photooxidative stress. Accordingly, ^1^O_2_ photoproduction, as measured from SOSG-EP fluorescence emission and PC-OH accumulation, was noticeably reduced in *OE:CYP81D11* relative to WT. In this context, it is interesting to note that, in wheat, another member of the CYP81D family, TaCYP81D5, was proposed to be involved in the response to salt stress through a possible modulation of ROS accumulation (Wang et al. 2020). Importantly, enhanced CO_2_ fixation and lowered ROS production in high light were found to provide a substantial advantage to *OE:CYP81D11* plants in terms of growth and biomass production.

On top of that, our RNAseq analysis also revealed the induction of various genes coding for protective mechanisms in plastids. In particular, *COR15A*, whose expression was markedly upregulated in response to *CYP81D11* overexpression, encodes a protein known to stabilize chloroplast membranes and protect cells against dehydration (Artus et al. 1996, Thalhammer et al. 2014). Partial folding of COR15 into α-helices upon stress-induced molecular crowding in the chloroplast stroma was reported to allow interaction of the protein with the non-bilayer MGDG lipid (monogalactosyldiacylglycerol) which in turn stabilizes membranes against hexagonal II phases (Thalhammer and Hincha 2014). Interaction between COR15 and MGDG may have a number of important functional implications because MGDG plays crucial roles in the structure and dynamics of chloroplast membranes, in particular under environmental stress conditions (Garab et al. 2016). Interestingly, increased expression was also found in *OE:CYP81D11* for genes encoding other COR proteins (*COR15B, KIN1 =COR6.6*) which are also involved in dehydration tolerance mechanisms (Wilhelm and Thomashow 1993, Guozhang et al. 2009, Thalhammer et al. 2014). Similarly, the chloroplastic lipocalin CHL/LCNP was induced in *OE:CYP81D11* relative to WT; this protein protects chloroplast lipids against oxidation under several abiotic stresses (Levesque-Tremblay et al. 2009, Malnoë et al. 2018). Considering their protective actions, it is clear that accumulation of plastid proteins coded by those genes can be very beneficial for chloroplasts under drought and photooxidative stress conditions.

The scheme of Fig. 8 summarizes our results and interpretations on the possible physiological roles of CYP81D11 in plant responses to environmental stresses. Irrespective of the exact molecular mechanisms underlying the action of CYP81D11, this study has identified this cytochrome P450 as a crucial component in the signaling pathway of the apocarotenoid β-CC leading to plant stress tolerance. An interesting aspect of the CYP81D11 function is its link with ^1^O_2_. In fact, β-CC is produced by ^1^O_2_ either by direct ^1^O_2_ oxidation of β-carotene or by induction of carotenoid cleavage enzymes such as LOX2 lipoxygenase (Havaux 2020). Accordingly, CYP81D11 is one of the most strongly induced genes by light in the Arabidopsis *ch1* mutant, which is an ^1^O_2_ overproducer (Ramel et al. 2013). Thus, induction of a gene by conditions associated with high ^1^O_2_ production ultimately leads to a decrease in ^1^O_2_ production, pointing to an interesting feedback regulation mechanism.

**Fig. 8.**
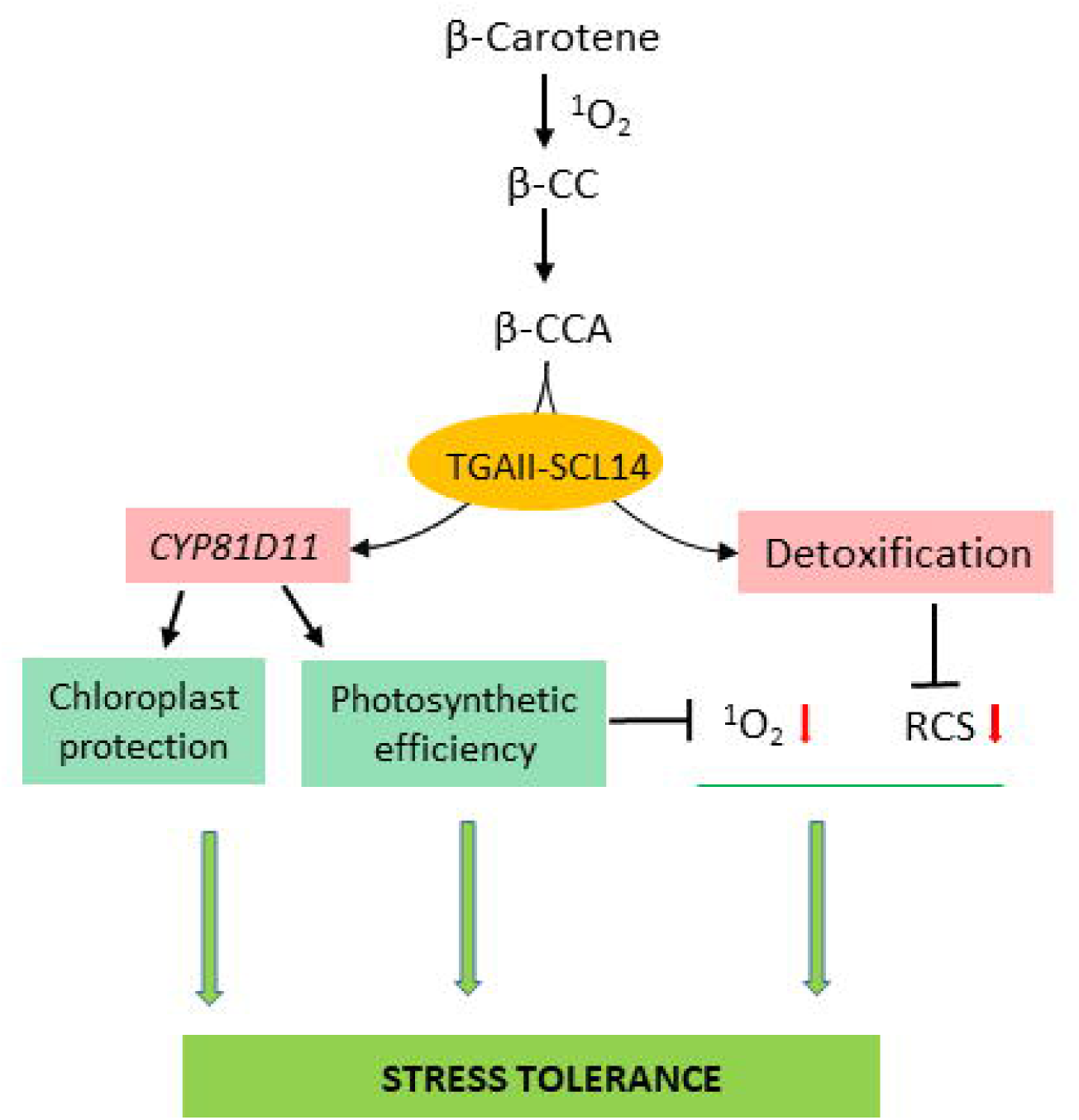
Schematic representation of the roles of CYP81D11 in plant stress responses. High CYP81D11 expression enhances the leaf photosynthetic capacity, thereby reducing ROS production in high light. High CYP81D11 levels are also associated with the upregulation of chloroplast protective mechanisms. These effects supplement the cellular detoxification pathway triggered by β-CC (D’Alessandro et al. 2018), boosting plant tolerance to pro-oxidative conditions, such as photooxidative stress.

Moreover, the enhancement of photosynthetic efficiency at high PFDs associated with the upregulation of CYP81D11 makes this gene a possible target for the improvement of photosynthesis in crop plants, especially in sunny environments.

## Methods

### Plant material and growth conditions

WT *Arabidopsis thaliana* (ecotype Col-0) was grown in potting soil in phytotrons of the Phytotec platform (BIAM, CEA/Cadarache) under controlled conditions of light (short day, 8/16 h (light/dark), PFD of 130 µmol photons m^−2^ s^−1^), temperature (21/18°C, day/night) and air humidity (ca. 60%). Several Arabidopsis mutant lines were used in this study: the triple mutant *tga256* (D’Alessandro et al. 2018), the *scl14* knockout mutant, a SCL14-overexpressing line *OE:SCL14* (Fode et al. 2008), *CYP81D11*- or *CYP72A13*-overexpressing lines (**OE:CYP81D11**, lines 5-5.3, 11-4.3, 12-2.1, and *OE:CYP72A13*, lines 3.2, 7.9, 5.2, respectively) (Bruce et al. 2008) and a *cyp81d11* knockout mutant (Matthes et al. 2010). The **OE:CYP81D11** lines were kindly provided by Christiane Gatz and Johnathan Napier. *OE:CYP72A13* lines were received from Johnathan Napier. The cyp81d11 mutant was obtained from NASC (SAIL_743_C10C1) and was screened for homozygosis by PCR.

In general, plants were grown for 5 weeks before treatments. Plants were exposed to photooxidative stress by transferring them to a growth chamber in which PFD was set at 1500 µmol m^-2^ s^-1^ and air temperature was 9°C (leading to an average leaf temperature of 18°C). Drought stress was imposed by stopping watering for 10 d (unless specified otherwise). The leaf relative water content RWC was calculated in % as 100 x (FW-DW)/(FWsat-DW) where FW is the fresh weight of leaf discs, FWsat the water-saturated weight (obtained by incubation of the leaf discs in water for 16 h) and DW the dry weight (obtained after leaf dehydration for 3 d in an oven at 90°C).

### β-CC and β-CCA treatments

β-CC was obtained from Sigma Aldrich or Astatech. Plants were exposed for 4 h to volatile β-CC in a transparent airtight box (≈22 L) in dim light (15 μmol photons m^−2^ s^−1^) at 22 °C, as previously described (Ramel et al. 2012b). 75 μL of β-CC were deposited on a wick of cotton to favor β-CC volatilization in the airtight box. For the control conditions, β-CC was replaced by distilled water.

For β-CC determination, leaf samples (around 200 mg fresh weight) were grinded in methanol containing 0.01% butylhydroxytoluene. After centrifugation, 5 µL of the supernatant was analyzed by LC-MS. A Kinetex C18 2.1⍰×⍰150⍰1mm 1.7⍰μm column (Phenomenex) was used with a flow rate of 300 µL min^-1^ achieved using a Vanquish UPLC system (Thermo Fisher, Waltham, MA, USA). A mobile phase A (water-acn 95/5, v/v) and mobile pahse B (acn-water (95:5, v/v), both containing 0.1% of formic acid, were utilized for elution. The elution was performed with a gradient: eluent B was increased from 5 to 95% in 26 min then maintained from 2 min, solvant B was decreased to 5% and then maintained for another 7 min for column re-equilibration. The column oven temperature was maintained at 35 °C. The effluent from the LC column was delivered to a Qexactive mass spectrometer operating in positive mode with the following conditions: sheath gas at 50; sweep gas at 2; auxiliary gas at 15; spray voltage at 3kV; capillary temperature at at 350°C; heater temperature at 400°C; S-lens RF at 45. β-CC identification was based on mass accuracy and MS/MS pattern. The extracted ion chromatogramm at 153.1274 with a tolerance of 5ppm was used for quantification based on a calibration curve obtained with pure β-CC standard solutions.

β-CCA was produced by oxidation of β-CC in water as previously described (D’Alessandro et al. 2019). β-CCA was applied to plants through the roots by watering the pots with a 1.5 mM solution of β-CCA (25 mL per plant). β-CCA was measured in leaves by GC-MS as previously described (Braat et al. 2023).

### Chlorophyll fluorescence

Chlorophyll fluorescence from attached leaves was measured with a PAM-2000 fluorometer (Walz). The maximal quantum yield of PSII photochemistry was measured in dark-adapted leaves by the Fv/Fm ratio where Fm is the maximal chlorophyll fluorescence level (measured with a 800-ms pulse of saturating white light) and Fv is the variable chlorophyll fluorescence (measured as Fm-Fo where Fo is the basal fluorescence level). The actual quantum yield of PSII photochemistry in the light (Φ_PSII_) was measured by calculating the (Fm’-Fs)/Fm’ ratio where Fm’ is the maximal fluorescence level and Fs is the steady-state level. NPQ was measured as 1-(Fm’/Fm). Plants were illuminated with different PFDs of white light provided by a Schott KL-1500 light source.

### Photosynthetic CO_2_ fixation and stomatal conductance

Gas exchange measurements were performed using a portable Li-Cor 6800 infrared gas analyzer (IRGA) (LI-COR Biosciences). A 6-cm² chamber was used to enclose the leaf, while the reference CO₂ concentration in the chamber was maintained at 400 μmol mol⁻¹. The flow rate was adjusted to 500 μmol s⁻¹, the chamber temperature was set to 23°C, and the leaf-to-air vapor pressure deficit (VPDleaf-air) was kept at 1.4 ± 0.5 kPa. Prior to measurements, the leaf was acclimated at a photosynthetically active photon flux density (PPFD) of 130 μmol m⁻² s⁻¹. After stabilization of the photosynthetic rate (A) and stomatal conductance (gsw) at this light level, the PPFD was increased to 1500 μmol m⁻² s⁻¹. The parameters A and gsw were continuously monitored and recorded until they reached steady-state levels.

### Photosynthetic pigments

Pigments were extracted in chilled 80% acetone buffered in 100 mM Tris-HCl pH 8. After centrifugation, the optical density of the supernatant was measured at 663.6 and 646.6 nm allowing the concentrations of chlorophyll a and chlorophyll b to be calculated as described in Chazaux et al. (2022).

### Clear-native polyacrylamide gel electrophoresis (CN-PAGE)

Thylakoids membranes were prepared from frozen shoots as described in Caffarri et al. (2009). Briefly, shoots were ground in 50 mL of B1 solution (hypertonic solution to preserve chloroplast structure, 20 mM Tricine/KOH, pH 7.8, 0.4 M sorbitol, 5 mM MgCl_2_, 10 mM NaF and the protease inhibitors 0.2 mM benzamidine, 1 mM ε-aminocaproic acid). After filtration with a nylon filter (mesh 30 µm) and centrifugation at 1400 g for 10 min at 4°C, the pellet was washed in B2 solution (isotonic solution, 20 mM Tricine/KOH, pH 7.8, 0.15 M sorbitol, 5 mM MgCl2, 10 mM NaF and 0.2 mM benzamidine, 1 mM ε-aminocaproic acid). After centrifugation at 4000 g for 15 min at 4°C, the pellet was resuspended in B3 solution (hypotonic solution to burst chloroplasts, 20 mM HEPES/KOH, pH 7.5, 15 mM NaCl, 5 mM MgCl2, 10 mM NaF). This solution was centrifuged at 4000 for 15 min at 4°C. Finally, the thylakoid pellet was resuspended in a small volume of B4 stocking solution (10 mM HEPES/KOH, pH 7.5, 0.4 M sorbitol, 15 mM NaCl, 5 mM MgCl2, 10 mM NaF), frozen in liquid nitrogen and stored at-80°C.

CN-PAGE was realized according to Järvi et al. (2011) with some modification as described below. Thylakoid membranes (17 μg of chlorophylls) were washed in 500 μL of 10 mM BisTris/HCl, pH 7, then centrifuged at 12000g for 10 min at 2°C. The pellet was resuspended in 25 mM BisTris/HCl pH 7 + 20% glycerol at 1 mg mL^-1^ chlorophyll final concentration. The same volume of detergent solution was added to obtain final concentrations of 0.5 mg mL^-1^ of chlorophyll in 0.9% (w/v) digitonin and 0.3% (w/v) n-dodecyl-α-D-maltoside (α-DDM). After 20 min of solubilization on ice in the dark, the solution was centrifuged at 12000 g for 10 min at 2°C. The solubilized complexes were loaded (17 μg of chlorophyll per lane) on a homemade clear-native polyacrylamide gels (CN-PAGE) using a 3.30–11% acrylamide:bisacrylamide (ratio of 29:1) gradient. The anode buffer was 50 mM BisTris/HCl pH 7 and the cathode buffer was 50 mM tricine +15mM BisTris/HCl pH 7 + 0.1% Deriphat and 0.02% α-DDM.

Fluorescence image of the gel was taken with a Fusion FX imaging system (Vilber) using a blue light transilluminator (centered at 470 nm) and a band-pass emission filter (690-720 nm).

Bands in the gel are indicated based on several previous papers using a similar system (for instance, Crepin et al. 2020).

### RNAseq

Total RNA was extracted from leaf samples by using Direct-zol RNA MiniPrep Plus (Zymo Research, R2072). Quantity and quality of RNA were assessed respectively using NanoDrop2000 (Thermo Scientific) and Qubit RNA IQ assay kit (Thermo Scientific). Samples were analyzed by BGI Genomics (Hong Kong), providing a stranded mRNA library, 20 M reads/sample, 100 bp paired-end reads (100PE) on DNBseq. Standard bioinformatics analyses were performed by the Doctor Tom platform of BGI Genomics.

### qRT-PCR

Total RNA was extracted using Direct-Zol RNA MiniPrep (Zymo Research) and treated with the RNase-free DNase Set (Zymo Research) according to the manufacturer’s instructions. Quantity of RNA was assessed using a NanoDrop2000 (Thermo Scientific). Reverse transcription was performed on 400 ng of total RNA using qScript cDNA SuperMix (Quantabio). Quantitative PCR was performed on a 480 Light Cycler thermocycler (Roche) using the manufacturer’s instructions, using SYBR Green I Master (Roche) with 10 µM primers and 0.125 µL of RT reaction product in a total volume of 5 µL per reaction. The qPCR program was: 95°C for 10 min, then 45 cycles of 95 °C for 10 s, 60 °C for 10 s and 72 °C for 10 s. The specific primers for PCR amplification, designed using the NCBI Primer designing tool, are listed in Supplemental Table S2. GAPC2 was used as reference gene for data normalization.

### Biochemical measure of lipid peroxidation

Lipid peroxidation was estimated by quantifying the hydroxy linolenic acid levels (HOTEs, hydroxyoctadecatrienoic acids) using the protocol of Montillet et al. (2004) as detailed in Ksas and Havaux (2022). Lipids were extracted from leaf samples (ca. 200 mg fresh weight) in methanol/chloroform, and HOTEs were quantified by HPLC-UV.

### Luminescence imaging of lipid peroxidation

A liquid nitrogen-cooled CCD camera (VersArray 1300B; Roper Scientific) was used to image the ultraweak luminescence associated with oxidative stress and lipid peroxidation in plants, as previously described (Birtic et al. 2011, Havaux and Ksas 2022). The back-illuminated CCD (CCD36-40; e2v Technologies) with a 1340⍰×⍰1300 imaging array operates at a temperature of −110°C. The sample was imaged on the sensor by a 50-mm focal distance lens with an f-number of 1.2 (F mount Nikkor 50-mm, f:1.2; Nikon) equipped with a shutter. Acquisition time was 20⍰min, and on-CCD binning of 2⍰×⍰2 was used to increase detection sensitivity, so that the resulting resolution was 650 x 670 pixels. Plants were dark-adapted for 2 h before measuring autoluminescence. Images were analyzed using Image J software.

### Reactive carbonyl species

Lipid carbonyls were measured in leaf samples (200⍰mg fresh weight) by HPLC-UV after derivatization with DNPH (dinitrophenylhydrazine). We used the protocol detailed in Mano and Biswas (2018) with a few minor modifications (Ksas et al. 2024): Addition of DNPH was done after centrifugation (7000*g* for 5⍰min), rather than decantation, and 50⍰µL (rather than 20⍰µL) were injected in the HPLC system. DNPH was obtained from Molekula, and carbonyl standards were obtained from Sigma-Aldrich and AccuStandard. Most of the standards were purchased in the derivatized form. The other standards were derivatized with DNPH as detailed in Mano and Biswas (2018).

### Singlet oxygen

The production of ^1^O_2_was measured in attached leaves from the fluorescence of the ^1^O_2_-specific singlet oxygen sensor green (SOSG) fluorescent probe (Invitrogen), as described previously (Ramel et al., 2013). With the help of a 1-ml syringe (without needle), 100 μM SOSG was pressure-infiltrated into the leaves through the lower surface. Plants with SOSG-infiltrated leaves were exposed to a PFD of 1500 μmol photons m^−2^ s^−1^ at 9 °C for 20 min. As a control treatment, plants with SOSG-infiltrated leaves were placed in the dark at room temperature for 20 min. SOSG-endoperoxide (EP) fluorescence was then measured from leaf discs punched from the SOSG-infiltrated leaves using a fiberoptics-equipped Perkin-Elmer spectrofluorometer (LS 50B) with a 475 nm excitation light. SOSG- EP fluorescence was measured at the peak emission (524 nm).

Hydroxy-plastochromanol-8 (PC-OH) is a specific marker of singlet oxygen in plants (Szymanska et al. 2014, Kruk and Szymanska 2021). PC-OH was extracted by grinding leaf samples (∼50 mg) in ethyl acetate. After centrifugation, the supernatant was filtered and evaporated on ice under a stream of N_2_. The residue was recovered in methanol/hexane (17/1) and analyzed by HPLC, as described elsewhere (Ksas et al. 2018). The column was a Phenomenex Kinetex 2.6 μm, 100 × 4.6 mm, 100 Å. The HPLC analysis was performed in the isocratic mode with methanol/hexane (17/1) as a solvent system and with a flow rate of 0.8 mL min^−1^. Hydroxy-plastochromanol-8 was detected by its fluorescence at 330 nm with an excitation at 290 nm. The standard was a kind gift from Dr. J. Kruk (Krakow, Poland).

## Acknowledgments

We would like to thank Christiane Gatz (Univ. Göttingen, Germany) and Johnathan Napier (Rothamsted Research, Harpenden, United Kingdom) for the kind gift of seeds. cyp81d11 seeds were obtained from the NASC Arabidopsis seed bank. Thanks are also due to the members of the Phytotec platform (CEA/Cadarache) for their help in growing plants under various environmental conditions. This work was financially supported by the French National Research Agency ANR (ApoStress project, ANR-21-CE20-0014) and by EDF-CEA GGP ‘Life Science’ 2023-2025.

## Author contributions

M.T. and M.H. designed the experiments and performed data analysis. M.T. performed most of the biological experiments. B.K. performed mst of the biochemical analyses (HOTE, PC-OH, RCS). M.T. performed the bioinformatic analysis. B.L. performed LC-MS analyses of apocarotenoids. S.C. performed the biochemical analysis of the photosynthetic complexes. M.H. and M.T. wrote the manuscript. All authors read and approved the final manuscript.

## Competing interests

The authors declare no competing interests.

## Additional information

**Supplementary information.** The online vesrion contains supplementary material: Figs. S1 to S9.

Tables S1 qnd S2.

## Notes

### Competing Interest Statement

The authors have declared no competing interest.

