## Supplemental Figures for "A cytochrome P450 involved in apocarotenoid signaling enhances plant photosynthetic capacity and photooxidative stress tolerance"

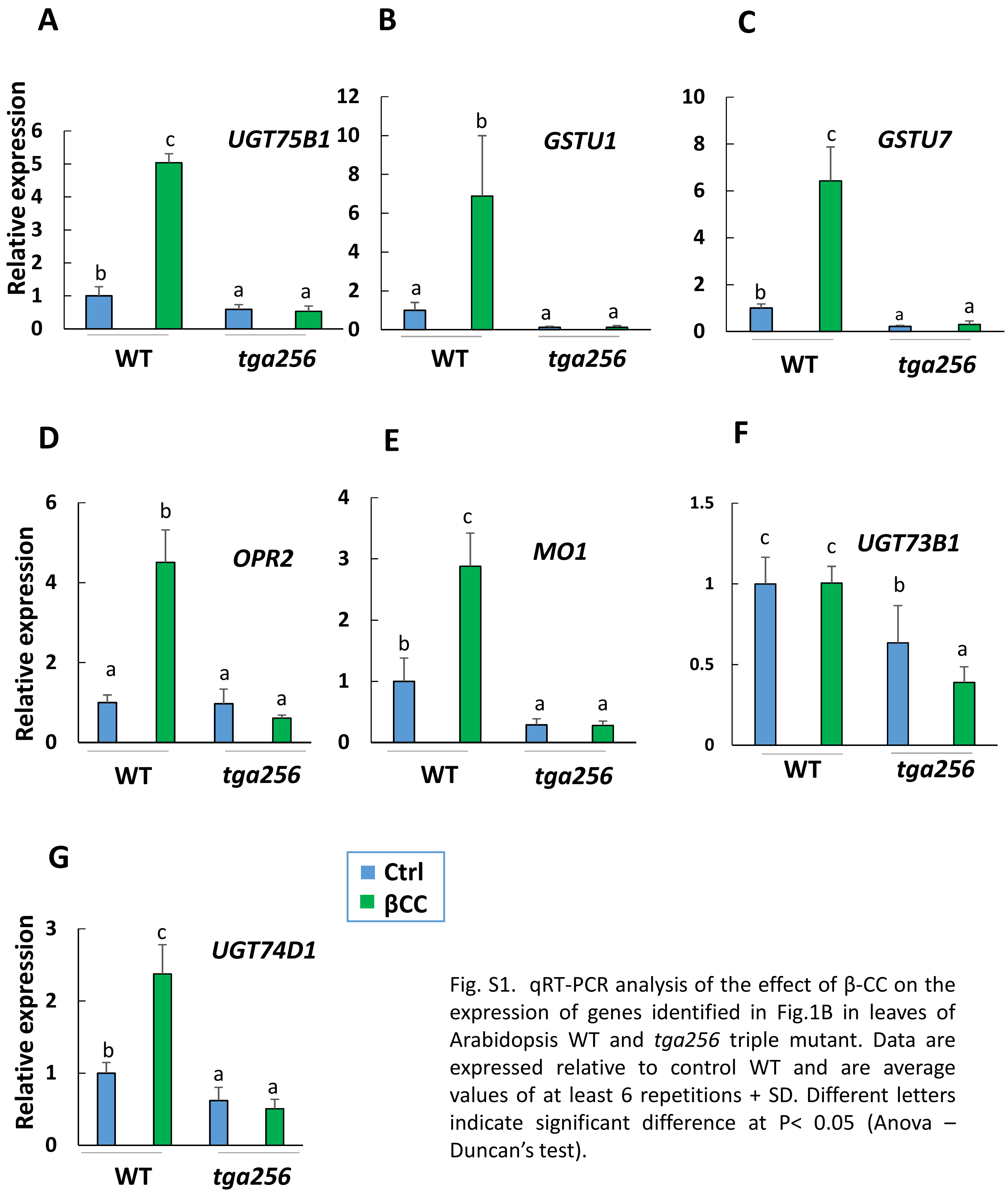


Fig. S1. qRT-PCR analysis of the effect of β-CC on the expression of genes identified in Fig.1B in leaves of Arabidopsis WT and *tga256* triple mutant. Data are expressed relative to control WT and are average values of at least 6 repetitions + SD. Different letters indicate significant difference at $P < 0.05$ (Anova – Duncan's test).

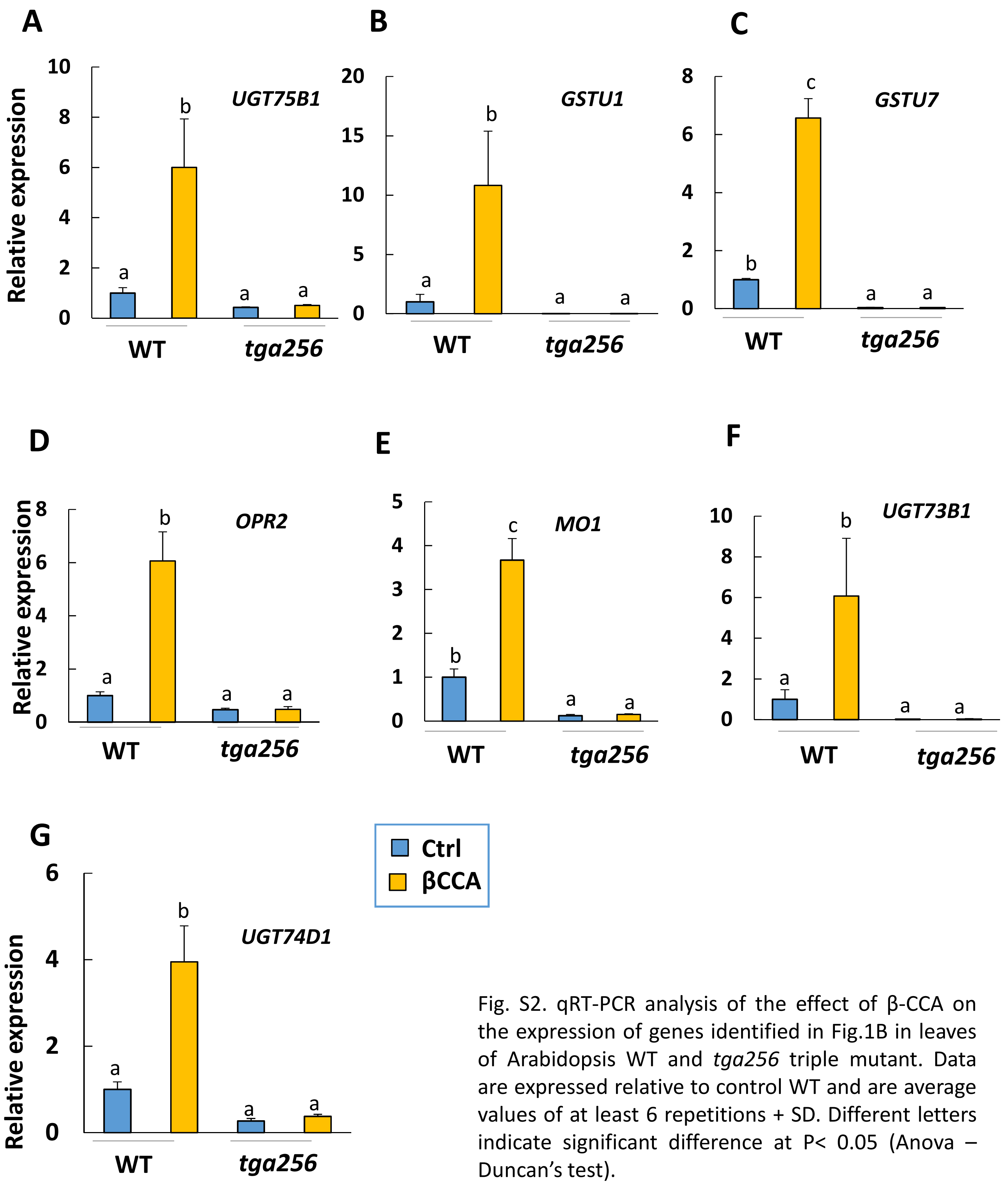


Fig. S2. qRT-PCR analysis of the effect of β-CCA on the expression of genes identified in Fig.1B in leaves of Arabidopsis WT and *tga256* triple mutant. Data are expressed relative to control WT and are average values of at least 6 repetitions + SD. Different letters indicate significant difference at P< 0.05 (Anova – Duncan's test).

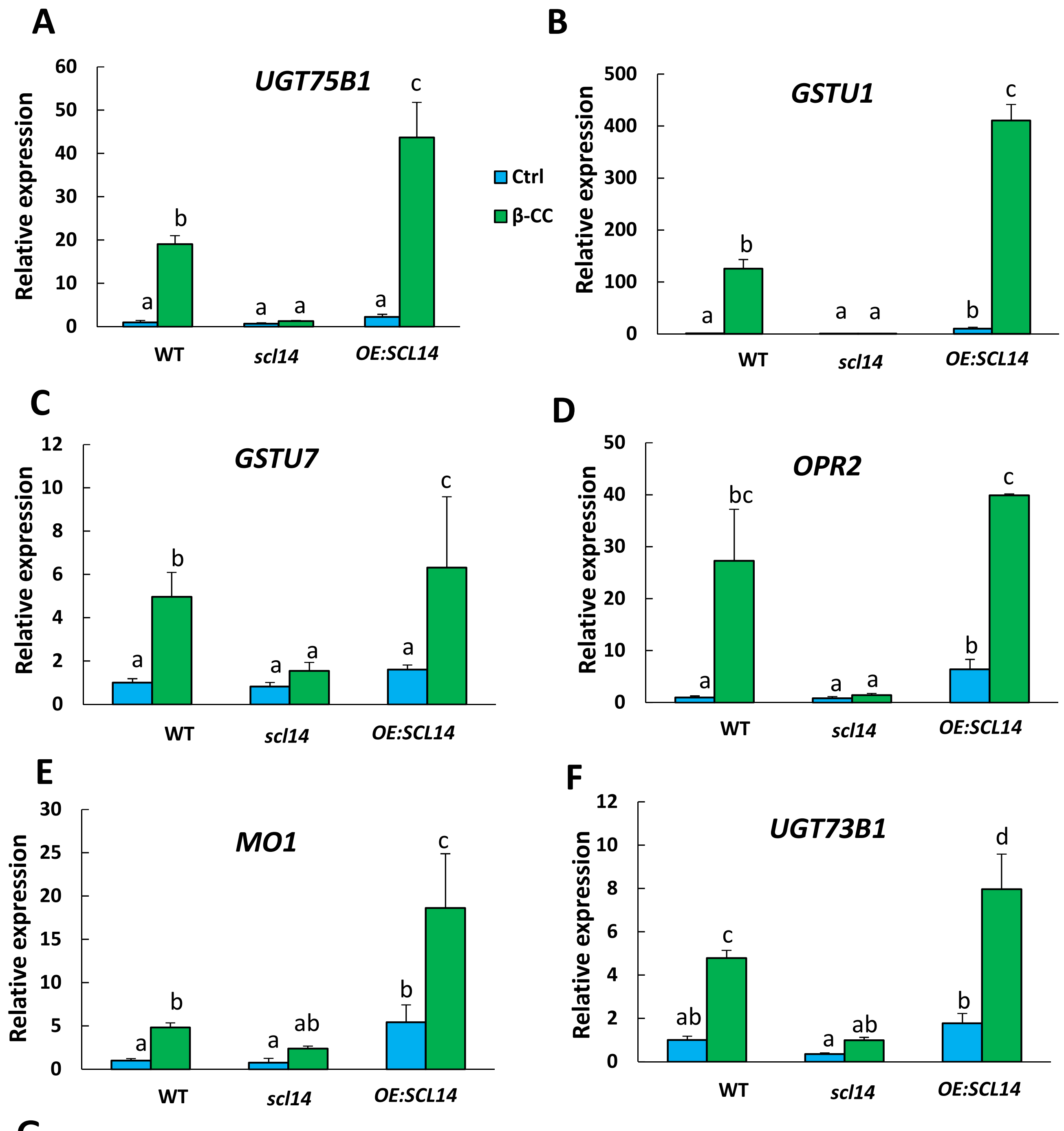


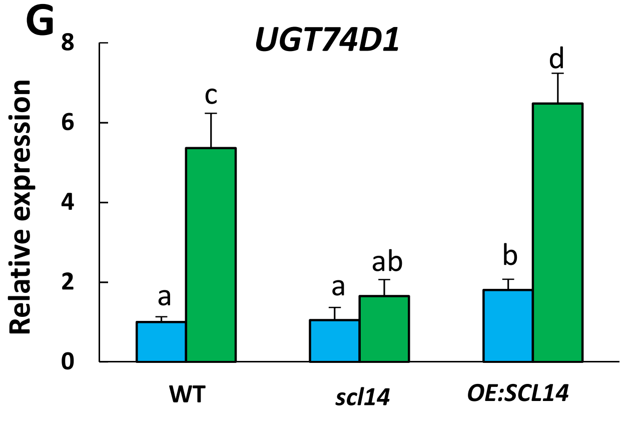


Fig. S3. qRT-PCR analysis of the effect of β-CC on the expression of genes identified in Fig.1B in leaves of Arabidopsis WT, *scl14* knockout mutant and *SCL14*-overexpressing line (*OE:SCL14*). Data are expressed relative to control WT and are average values of at least 6 repetitions + SD. Different letters indicate significant difference at P< 0.05 (Anova – Duncan's test).

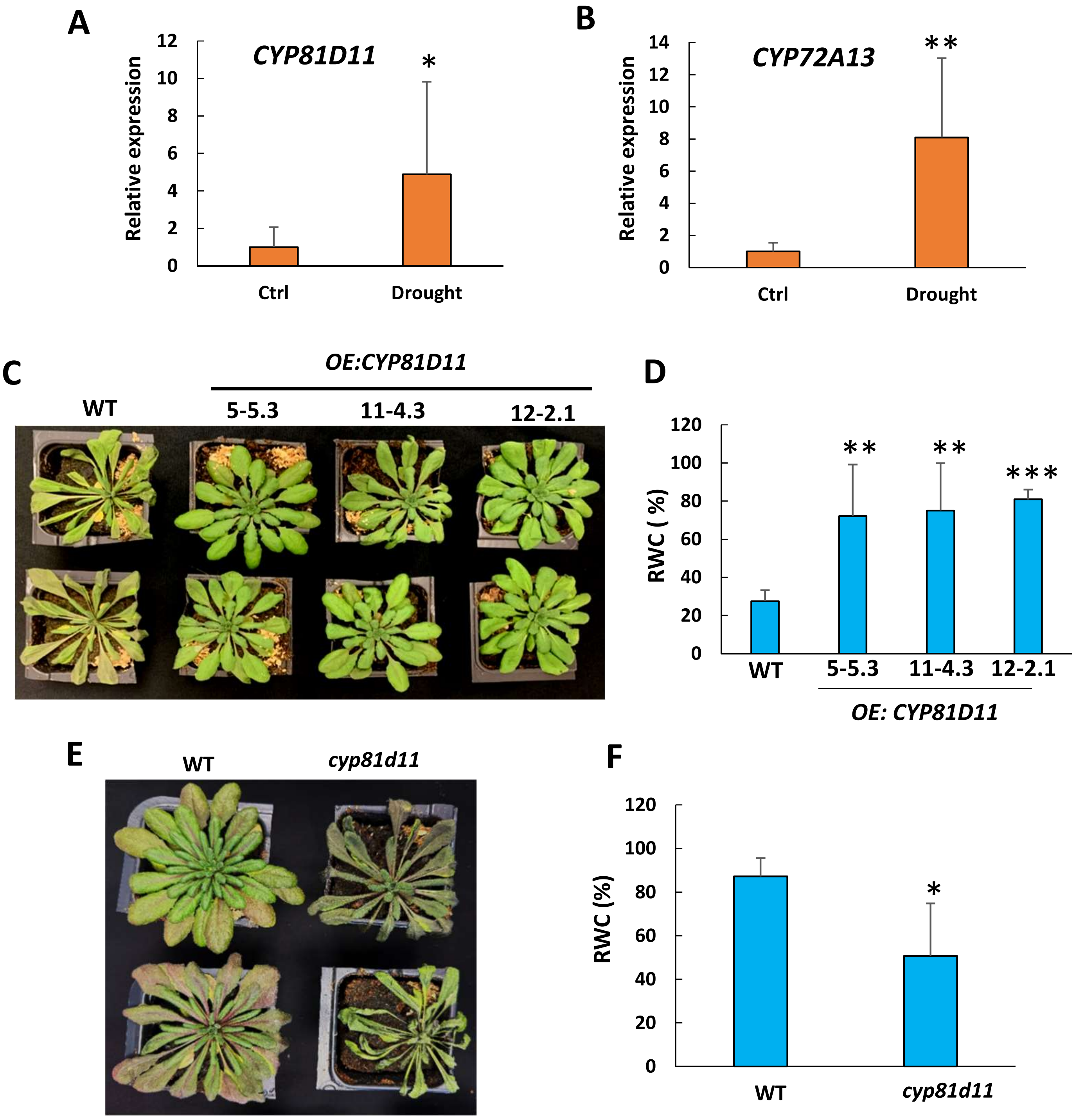


Fig. S4. Effects of drought stress on Arabidopsis WT, *OE:CYP81D11* and *cyp81d11* mutant plants. Water stress was imposed by stopping watering (8 d for panels A-B nd E-F, 10 d for panels C-D). A-B) qRT-PCR analysis of *CYP81D11* and *CYP72A13* gene expression in response to drought stress. Data are expressed relative to control WT and are average values of at least 6 repetitions + SD. C) Picture of water-stressed WT and *OE:CYP81D11* plants (3 lines, 5-5.3, 11-4.3 and 12-2.1). D) Leaf RWC of water-stressed WT and *OE:CYP81D11* plants. Data are average values of 10 repetitions + SD. E) Picture of water-stressed WT and *cyp81d11* mutant plants. F) Leaf RWC of water-stressed WT and *cyp81d11* mutant plants. Data are average values of 15 repetitions + SD. ***, ** and *, significant difference at P< 0.001, P< 0.01 and P> 0.05, respectively (Student's t-test).

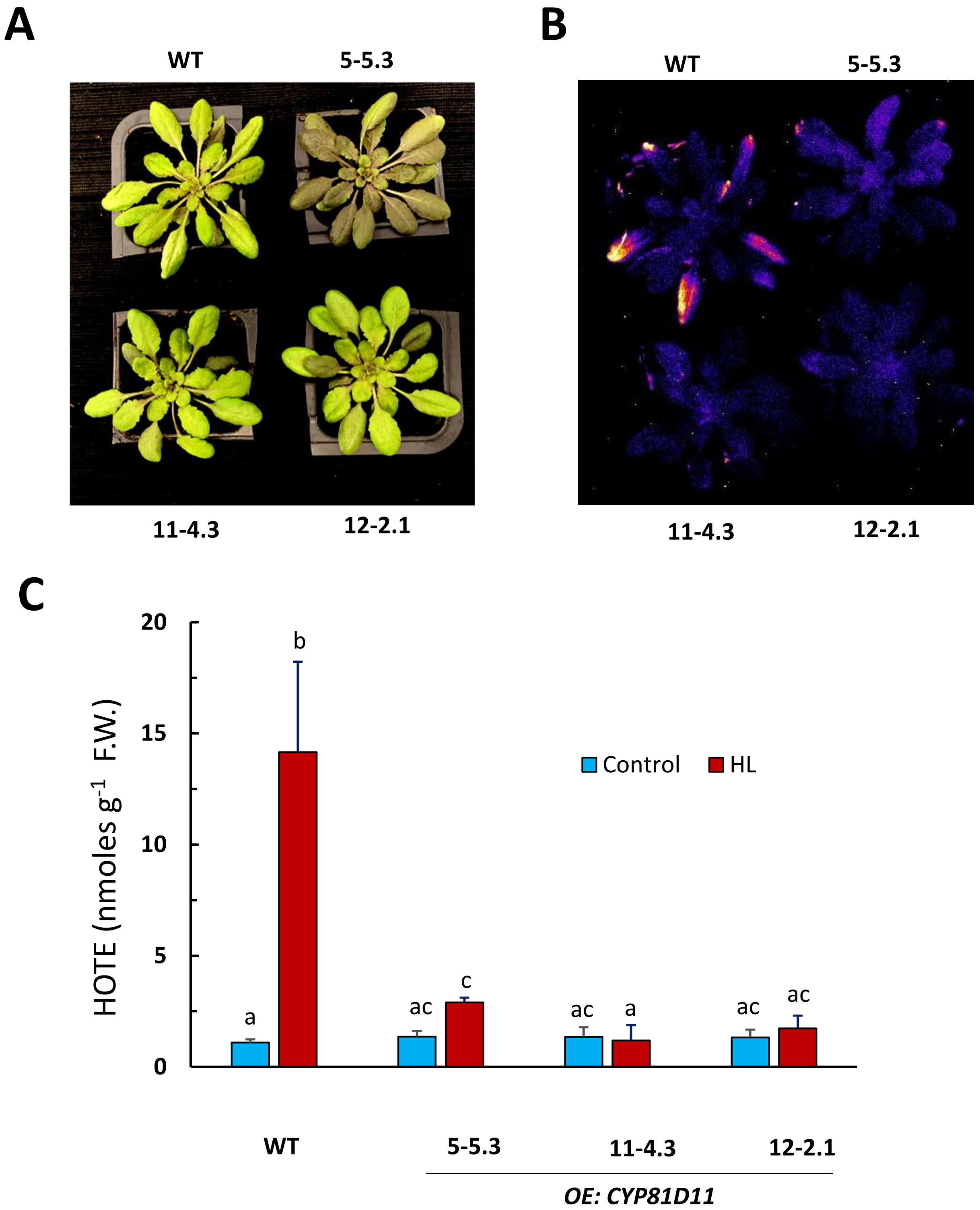


Fig. S5. Effect of high light stress on several Arabidopsis lines overexpressing the *CYP81D11* gene (*OE:CYP81D11*, lines 5-5.3, 11-4.3 and 12-2.1). Plants were exposed for 2 d to 1500 µmol photons $m^{-2}$ $s^{-1}$ at 9°C air temperature. A) Picture of the plants after high light stress. B) Autoluminescence imaging of lipid peroxidation after high light stress. C) HOTE levels in leaves before and after high light stress. Data are mean values of at least 3 repetitions + SD. Different letters indicate significant difference at P< 0.05 (Anova – Duncan's test).

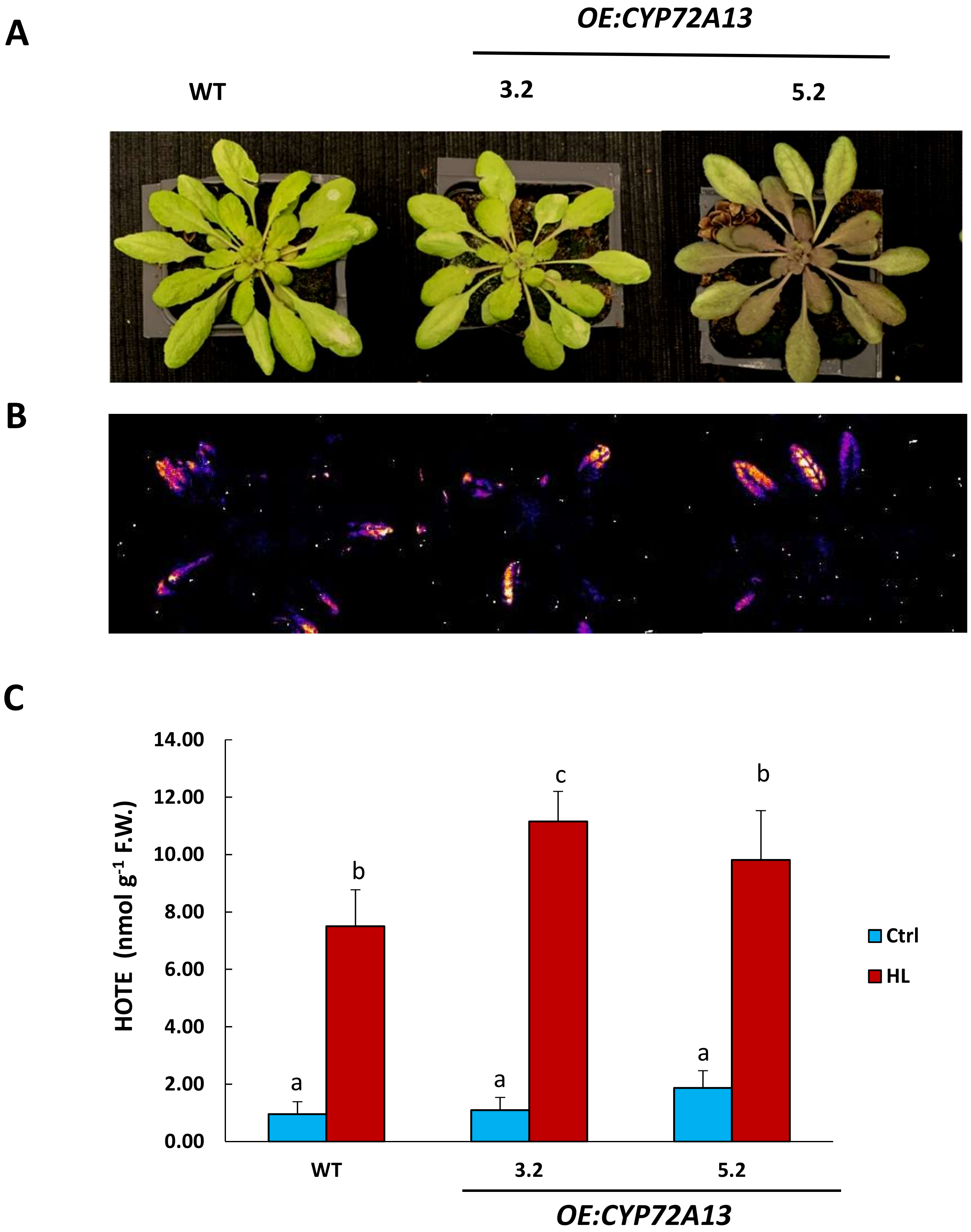


Fig. S6. Effect of high light stress on two Arabidopsis lines overexpressing the *CYP72A13* gene (*OE:CYP72A13*, lines 3.2 and 5.2). Plants were exposed for 2 d to 1500 µmol photons $m^{-2}$ $s^{-1}$ at 9°C air temperature. A) Picture of the plants after high light stress. B) Autoluminescence imaging of lipid peroxidation after high light stress. C) HOTE levels in leaves before and after high light stress. Data are average values of at least 3 repetitions + SD (Anova – Duncan's test).

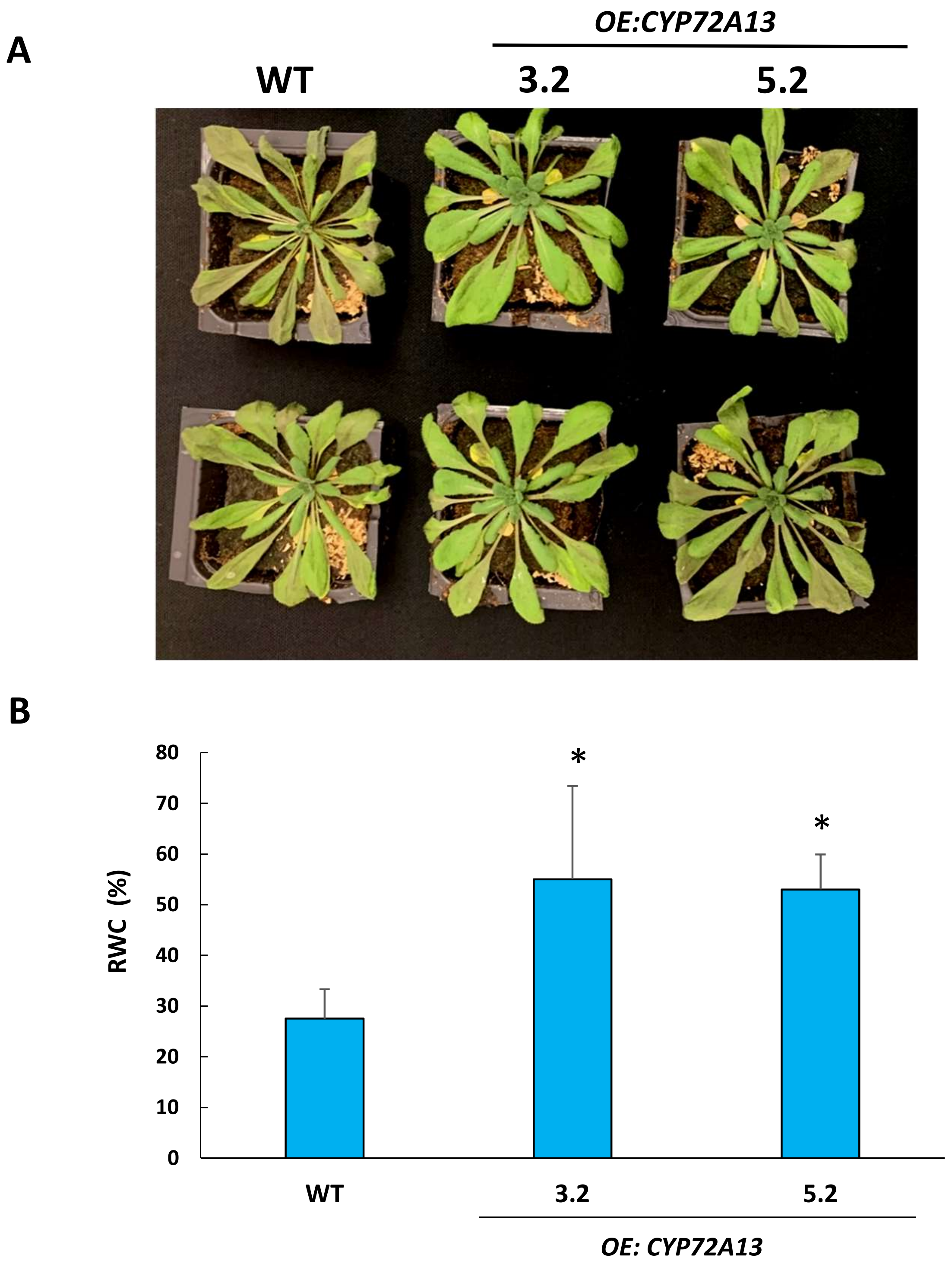


Fig. S7. Effect of drought stress on two Arabidopsis lines overexpressing the *CYP72A13* gene (*OE:CYP72A13*, lines 3.2 and 5.2). Plants were exposed to water stress by stopping watering for 10 d. A) Picture of the plants after drought stress. B) Leaf RWC after drought stress. Data are average values of 10 repetitions + SD. *, significant difference at $P < 0.05$ (Student's t-test).

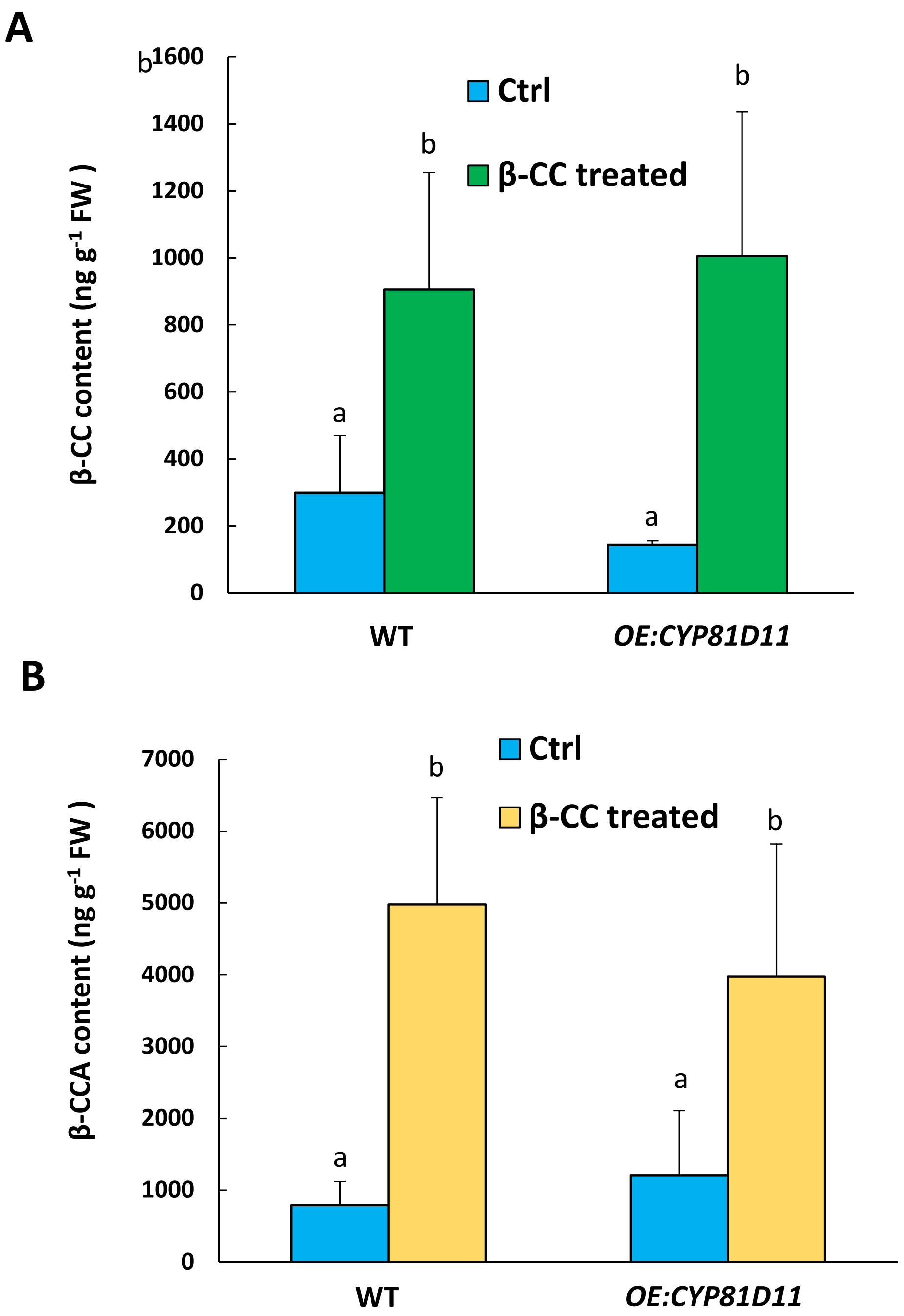


Fig. S8. β-CC and β-CCA content in leaves of Arabidopsis WT and *OE:CYP81D11* plants before and after 4-h exposure to volatile β-CC. A) β-CC content. B) β-CCA content. Data are average values of 3 repetitions + SD (Anova – Duncan"s test).

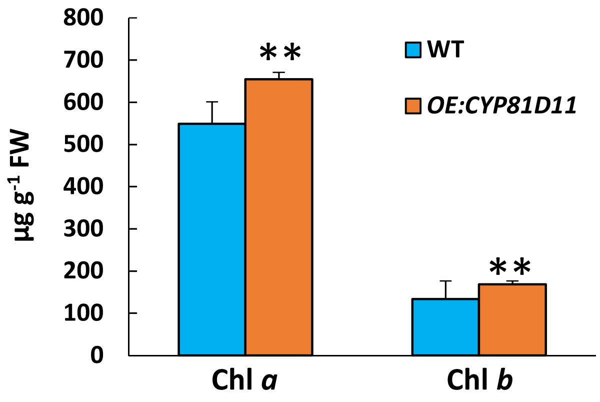


Fig. S9. Chlorophyll *a* and chlorophyll *b* content in Arabidopsis WT and *OE:CYP81D11* leaves. Data are average values of 6 repetitions + SD. **, significant difference at P< 0.01 (Student's t-test).
