## Supplemental Tables for "A cytochrome P450 involved in apocarotenoid signaling enhances plant photosynthetic capacity and photooxidative stress tolerance"

**Table S1: Analysis of TGACG binding motif:** PROMOTER ANALYSIS OF UPSTREAM 600BP FROM +1 SITE

| Gene | Motif Sequence | Position | Strand | Hit Sequence |
|---|---|---|---|---|
| ***CYP81D11*** | TGACG | 231 | - | TGACG |
| ***UGT75B1*** | TGACG | 196<br>208 | +<br>+ | TGACG |
| ***GSTU1*** | TGACG | 242<br>254 | -<br>- | TGACG |
| ***GSTU7*** | TGACG | 76<br>79<br>88 | +<br>-<br>+ | TGACG |
| ***OPR2*** | TGACG | 535<br>544 | -<br>+ | TGACG |
| ***MO1*** | TGACG | 163<br>176 | +<br>+ | TGACG |
| ***CYP72A13*** | TGACG | 312 | - | TGACG |
| ***UGT73B1*** | TGACG | 517<br>529 | -<br>- | TGACG |
| ***UGT74D1*** | TGACG | 157 | + | TGACG |
| ***AT4G01330*** | TGACG | 507 | + | TGACG |

### Table S2: Primers used during this study

| | qRT-PCR primers | Forward | Reverse |
|---|---|---|---|
| Toll like interlukin | AT3G50970 | AAGCTTCCCGGTGGTCATC | GCGACTCAATGAAAGAAAGCC |
| GSTU5 | AT2G29450 | TACTTCGGAGGCAAGACTGT | CCTTCCCAACCTCTCTCCAA |
| MAPKKK18 | At1g05100 | AGTGATATATGGGCGGTGGG | GGTAAAATCCGCCCCAATCC |
| SDR1 | AT4G13180 | AAGCAGATGTGTCAGACCCG | ACCTGCACAATTCACGACGA |
| AER | AT5G16970 | CCATGGTAAGAATGTTGGGAAACAA | AAAACAAAAACCACCCACACAAACT |
| ANAC 102 | AT5G63790 | AGCTTCCAGAAATGGCGTTGT | AATAACCGGTTCCAGCTGCC |
| UGT75B1 | AT1G05560 | AGGAGGTGTTTGGAAGCCGT | CTCTACCCGCTTCCATCGCT |
| GSTU1 | AT2G29490 | TGCCCAACAAGACCCCTTTG | TTGTGGACAAGAACCGGAACTT |
| GSTU7 | AT2G29420 | TGTGACGGCGATGAAAGTTGT | ATCTCTCGTCGCTTCAACCAC |
| OPR2 | AT1G76690 | ACGGGGAAGCCAATTATGCC | CGTCTTGGAGGGGTAAAGCG |
| MO1 | AT4G15760 | GATTGGATGCGATGGTGCCA | TCACCGCACGACAAGCAAAC |
| CYP72A13 | AT3G14660 | AACCCGGCTTTCCACCTTGA | TTCGCCAACAATCTCGCTGC |
| UGT73B1 | AT4G34138 | GCTTTGGCGTGTTGGTGAAC | GGACCGATATGCCACGCTCT |
| UGT74D1 | AT2G31750 | GCCCAAGTCAACGAATGCCT | AGACGGCCAAGCTTCCAAAA |
| CYP81D11 | AT3G28740 | CGACGATCTTGCCCTGGTTC | TCTTCAACCCTCTCCCACTCA |
| GAPC2 | At1g13440 | GGCCATCAAGGAGGAATCTG | CTTGGCATCGAAAATGCTTG |
| | Genotyping Primer | | |
| | AT3G28740 | GAACAAAATAAACCGATGGCC | CGCTAGACAAATTATGAGCCG |
